# Dominance hierarchies are structured similarly in females and males across primate groups

**DOI:** 10.64898/2026.08.14.744899

**Authors:** Lucas Spicher, Elise Huchard, Dieter Lukas

## Abstract

Classic socio-ecological theory predicts that males and females experience different sources and mechanisms of social competition. Whether these differences translate into sex-specific structural properties of dominance hierarchies remains unclear. Here, we compiled 156 dominance interaction matrices from 80 published studies and extracted three commonly used metrics - hierarchy steepness, linearity and the directional consistency index - to investigate the structural characteristics of male and female dominance hierarchies across primates. All three metrics were strongly affected by methodological and demographic variables. Steepness increased with the number of recorded interactions and group size, linearity decreased as matrices became sparser, and directional consistency declined with increasing numbers of interactions. Steepness covaried positively with both linearity and directional consistency, indicating that groups with steeper hierarchies also exhibited more linear and more directionally consistent relationships. We found no sex differences in steepness, linearity or directional consistency. These results suggest that current metrics primarily reflect variation in the sampling effort and the rate of interaction of the recorded behaviour and appear therefore not to capture potential sex differences in the forms of competition. Our findings highlight the need for alternative measures of power asymmetries that are less confounded by sampling effort and demographic variation to better understand how competition and conflict are structured across primate societies.

## 2. Introduction

In mammals, dominance hierarchies are a core part of social structure, particularly in species that live in social groups. Hierarchies represent a pattern of repeated, agonistic interactions between individuals (1), that shape access to resources and directly influence survival and fitness (2,3). Given that sexes often compete over different resources and via different mechanisms (4–6), it has often been assumed that their dominance hierarchies may differ in their form and properties (7). Adult male mammals are frequently involved in aggressive interactions, such as biting, hitting, or chasing, when competing over access to mates. These interactions among the males can lead to serious injury (8), which can be associated with frequent rank changes (9). In contrast, females may be less likely to resolve contests through direct physical aggression given that competition may provide less benefits while carrying risks for dependent offspring. Accordingly, female hierarchies may be more signal-based, with dominance relationships maintained through ritualized signals such as supplants, threat faces, or avoidance behaviour (10,11). Additionally, where females are philopatric as in many mammals, female ranks are often established early and could be more stable due to kin-biased support and the longer reproductive lifespan of females (7,10). Despite these general assumptions about potential sex differences in conflict resolution across mammals, we still lack a clear understanding of whether, and if so how, these sex differences link to differences in the structural properties of dominance hierarchies.

Dominance structures vary extensively across species, likely reflecting ecological contexts and social conditions (12,13). In particular, primatologists have distinguished different types of societies, ranging from so-called despotic systems, characterized by steep and linear hierarchies, to more egalitarian systems, marked by weaker and more relaxed hierarchies (12–15). For example, female rhesus macaques (*Macaca mulatta*) manage conflict through strict rules of submission, exhibit steep hierarchies and rarely reconcile conflicts, while female Tonkean macaques (*Macaca tonkeana*) exhibit shallower hierarchies, often counter-attack after being aggressed, and resolve conflicts through frequent reconciliations (16). Across macaque species, this covariation among social traits related to dominance and conflict mitigation has been formalized under the ‘social style’ or ‘dominance style’ frameworks (12,14,16). Though interspecific variation in social style is well documented, far less is known about how males and females differ in these behavioural syndromes. In this study, we build on these empirical observations to determine whether there are systematic sex-differences in dominance hierarchies.

A key property to characterize structural variation in dominance hierarchies has been steepness—formerly known as dominance gradient – which measures individuals’ success rate at gaining fights, in relation to their ordinal rank, which reflects their position in the hierarchy (17). The steeper a hierarchy is, the higher is the probability that a high-ranking individual will win an agonistic interaction against a subordinate. The most despotic macaque species (14,16) (e.g. grade 1: *Macaca mulatta*) exhibit higher steepness values than the most tolerant species (e.g. grade 4: *Macaca tonkeana*) (18) suggesting a link between social style and hierarchical steepness. In chimpanzees, variation in steepness is accompanied by other changes in social behaviour, but these do not exactly match the macaque social styles: in steep hierarchies, males use grooming as a bargaining tool to gain support in conflicts, whereas in shallow hierarchies, this strategy is not observed (19). Likewise, a comparative analysis of six captive bonobo groups found that variation in hierarchy steepness was associated with variable grooming patterns (20). Overall, variation in steepness may reflect critical aspects of dominance hierarchies, so that understanding such variation may open broader perspectives on the architecture of social structure across populations and species.

Other properties of dominance hierarchies include linearity and sparseness. Linearity (generally reflected by Landau 1951 h’) measures how well dominance relationships form a transitive, complete order: in linear hierarchies, all dyads are decided (one individual dominates the other rather than there being a tie) and triads are transitive (if A > B and B > C, then A > C) (21). Linearity depends on both the number of established relationships and their transitivity, while steepness quantifies absolute differences in dominance success between adjacent ranks using cardinal measures; the two thus are expected to capture complementary aspects—structural completeness vs. rank disparity (17). Sparseness reflects the proportion of unknown/undecided dyads in the interaction matrix, arising from limited observations or true non-interactions (22). Higher sparseness reduces both linearity and steepness by limiting decided dyads, often yielding shallower, less consistent hierarchies (22,23). The Directional Consistency Index (DCI) has been proposed as an alternative that is less sensitive to this potential methodological limation. The DCI quantifies the degree to which interactions within dyads are consistently directed in the same direction, ranging from 0 (bidirectional) to 1 (completely unidirectional) (24). However, the potential advantage of this alternative approach should be regarded not as absolute given the lack of a general demonstration of its resistance to the effects of sparseness (25).

The steepness, linearity, and DCI of dominance hierarchies could be influenced by sex differences in the sources and mechanisms of intrasexual competition. In addition, the hierarchies of females and males might also differ because of socio-demographic factors such as group composition. Group size can influence steepness and linearity by affecting the proportion of unknown relationships and matrix completeness, with larger groups often yielding more unknowns and inconsistent rank orders (22). Because primate groups usually have a larger number of females than males, female hierarchies might become less linear and stable as group size increases (26). Finally, power asymmetries between the sexes, which vary across societies (27) could also, potentially, influence the structure of their hierarchies. For example, the dominant sex may exhibit a steeper hierarchy simply because dominant individuals engage more readily in agonistic interactions. Beyond this, sex differences in the relative use of aggressive versus submissive signals — and more broadly in which sex socially dominates the other — could further shape the structure of dominance hierarchies (28).

To date, evidence for sex differences in hierarchical steepness or other measures of hierarchical structure remains limited. In bonobos, where females dominate males socially, hierarchical steepness was found to be higher in males than in females (29). Similarly, in the despotic societies of rhesus macaques, males were found to exhibit higher steepness than females (30). One comparative analysis across mammals found no overall effect of sex on hierarchical steepness, but agonistic interactions were more directionally consistent in male-only than in female-only groups (31). However, this analysis included groups of all ages, in both captive and naturalistic settings and did not specifically examine potential demographic drivers of sex differences in dominance structure. Another comparative analysis found that the relative prevalence of aggression and ritualized submission in both sexes was sensitive to variation in the degree of female dominance across species. In societies where female dominance prevails, individuals used more submissive signals and fewer aggressive acts (28). Notably, hierarchies inferred from different agonistic behaviours are not always comparable (32) suggesting that the choice of behaviour used to construct dominance matrices may itself contribute to observed sex differences.

Here we present a comparative analysis examining the extent and potential sources of male-female differences in hierarchical properties, namely steepness, linearity and DCI, using a large sample of studies from haplorrhine primates. This study has five main aims: (1) evaluate the robustness of our structural measures of dominance hierarchies (i.e, steepness, linearity and DCI) to differences in sampling and demography, namely sparseness, number of interactions, study duration and number of individuals forming the hierarchy, and the type of behaviour used to build dominance matrices (comparing signal-based and aggression-based hierarchies); (2) determine whether steepness, linearity and DCI covary across species, reflecting specific systems of dominance structures, (3) compare steepness, linearity and DCI in male versus female hierarchies, (4) test whether any sex difference in these properties may be linked to socio-demographic properties (group size, number of individuals, or number of interactions), or (5) to sex differences in agonistic behaviour, such as variation in which sex socially dominates the other or (6) in the nature of behaviours used during competition (agonistic acts versus signals).

## 3. Materials and Methods

### Dominance data collection

We used published dominance matrices, reflecting data of pairwise aggression, displacement, or submission behaviours, to examine the properties of hierarchies. We accessed these matrices from the open database of the EloSteepness.data R package (33). The database consists of 610 dominance matrices collected from scientific publications (articles, reviews, doctoral and master theses). Our extraction and preparation of dominance data from the database was conducted from January 2025 to August 2025. We focused on matrices of primate species, of which there are 271 in this dataset. For each matrix, we inspected the original publication (n=136 candidate publications) to check whether the individuals in the dominance matrix were clearly identifiable, by carrying out two checks. We first checked that the sex of each individual in the matrix was available. If the matrix consisted solely of females or males, this resulted in one data point in our sample of a dominance matrix from which we calculated the structural properties. If the matrix consisted of both males and females, we split the matrix into two sex-specific matrices, resulting in two data points in our sample. Studies that did not provide any information on the sex of the individuals were excluded. Second, we checked that information on individual ages or age classes was available. We only included behavioral data from adults in our analysis. When the matrix was composed of different age classes, we manually removed non-adults (infants, juveniles, and subadults). When the authors provided age information (numerical) and not all individuals were adults, we checked if the authors provided an age threshold for adulthood in their population. If so, we used their threshold to exclude non-adults. When no adulthood threshold was proposed, we used a reference database on ageing (*” AnAge: The Animal Ageing and Iongevity Database”:* https://genomics.senescence.info/species/index.html) to determine the age at sexual maturity of the relevant species. In cases where such information was not available for a specific species from these databases, we conducted an additional search on Google Scholar using the species name, potentially the location where hierarchical data was collected, and keywords such as “adulthood”. In the absence of any indication to differentiate adults from non-adults, the study was discarded from our sample. After applying these two criteria, we obtained 156 dominance matrices (99 for females and 57 for males), derived from 80 papers (Figure 1).

**Figure 1:**
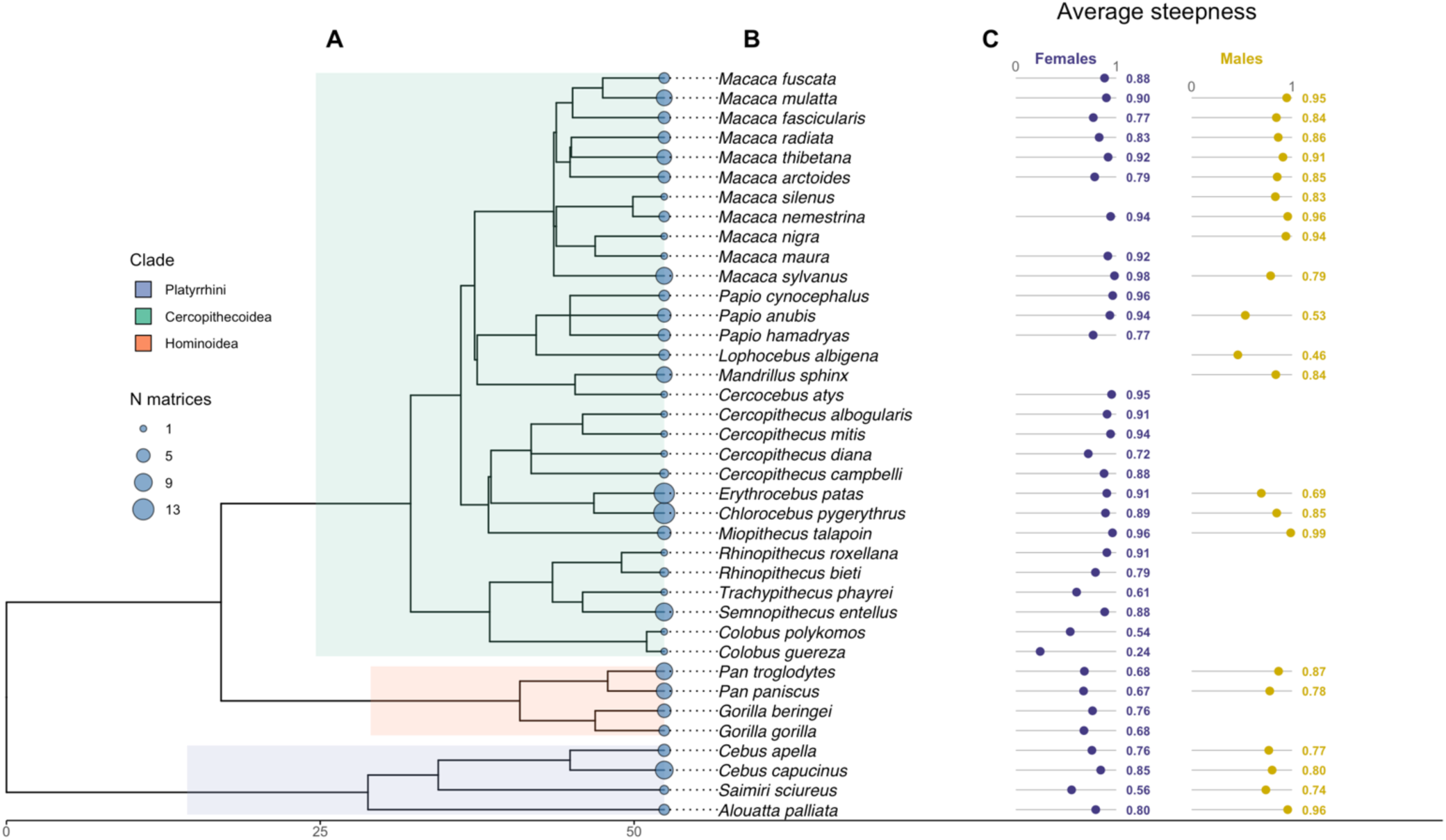
For the primate species in our dataset, the (A) phylogenetic relatedness, (B) number of dominance matrices available, and (C) the average steepness of matrices for males and females. Each tip represents a species (Latin name listed in italics), and the point size at the tip indicates the number of dominance matrices included for that species in the dataset. In our dataset, we have data on the steepness of hierarchies of males (shown in gold) for 21 species, and for females (shown in purple) for 34 species.

### Data on study features

For each matrix, we scored a set of descriptive metrics: the number of same-sex individuals in the group, the total number of interactions in the matrix, and the proportion of unknown relationships in the matrix (i.e., dyads with no observed interactions, known as “sparseness”). Whenever possible, we also collected data on the period over which the data were collected, the type of interactions recorded (aggressive acts, submission signals, both) and if the subjects in the study were wild or captive. We distinguished aggressive acts from signals following Kappeler et al (2022) (28): “Structurally, we distinguished between acts, which involve physical contact or locomotion, such as lunging or fleeing, and visual or vocal signals, such as non-physical threats or grimacing”. To test whether steepness reflects which sex dominates the other, we collected data on intersexual dominance from Huchard et al. (2025) (27), who provide, for a set of primate species, the proportion of intersexual agonistic interactions won by females. This variable was available for a subset of our sample (n = 141 matrices from 27 species) and was used as a continuous predictor of steepness in a dedicated model (see Statistical analyses).

### Steepness, linearity and DCI values

Steepness is calculated as the absolute slope of the regression of individuals’ cardinal dominance scores — numerical values obtained according to the chosen dominance index — on their ordinal ranks, which are defined as the ascending order of these scores (17). To calculate steepness, we relied on the Bayesian Elo-rating approach, which provides a statistical framework that accounts for uncertainty in dominance interactions and allows for more robust comparisons between groups and species (23). The linearity of a hierarchy is a structural property that reflects the degree to which dominance relationships can be ranked in a linear order (21). The standard measure of linearity, the h’ index, reflects the proportion of pairwise relationships in the group that are transitive, i.e., that are in the ranking order that minimizes hierarchical inconsistencies. The linearity of complete matrices (without missing data, with sparseness of 0) was characterized by the h index, while for incomplete matrices, with interactions containing zeros, we used the adjusted h’ index (21). The Directional Consistency Index (DCI) was extracted for each group as a measure of the asymmetry of dyadic dominance relationships. DCI values range from 0 (interactions within dyads are on average evenly balanced, i.e. no directional asymmetry) to 1 (all interactions consistently directed in the same direction within dyads). All three measures (steepness, linearity and DCI) were bounded between 0 and 1.

We calculated the median steepness of each matrix with Elo-ratings in a Bayesian framework (23), in the statistical software R version 4.5.2 using the function “elo_steepness_from_matrix” from the R package “EloSteepness” (34). We ran 2,000 iterations with a “warm-up” of 1,000 iterations over 4 MCMC chains generating 4,000 posterior samples. Linearity was determined using the “h.index” function from the “EloRating” R package (35). Finally, the DCI was estimated using the “dci” function from the “compete” R package (36).

### Information on Phylogenetic Relatedness

We used a time-calibrated mammalian supertree (37) to reflect the phylogenetic relationships among species in our sample. We matched the species names to those in the tree, and trimmed the tree using functions of the package ‘ape’ (38) in R.

## Statistical Analyses

We performed all analyses in the statistical software R version 4.5.2 (39). We provide the code at https://github.com/dieterlukas/DominanceHierarchyDifferences (40).

### Estimation of Phylogenetic Signal

Pagel’s λ (41) and Blomberg’s K statistics (42) were used to assess phylogenetic signal separately for male and female hierarchies. Closely related species were tested to see if they exhibited more similar steepness and linearity values than expected by chance. We estimated phylogenetic signals and assessed their significance using randomisation tests implemented with the’phylosig’ function in the package ‘phytools’ (43).

### Estimation of Associations Among Variables

We used Bayesian models implemented with the R package “rethinking” (44) and Stan via MCMC (45) to estimate the associations between properties of the hierarchy and the predictors. In all analyses, we set broad priors centered on zero (reflecting, a priori, no relationship). We run all models across four chains with 1000 iterations each drawing samples from the last 500. For all estimated relationships, the diagnostic criteria indicated that the chains were efficient (number of effective samples larger than 500) and had converged (Gelman-Rubin convergence diagnostic, R-hat, <1.01).

For the first objective, to evaluate the robustness of our hierarchy measures, we fitted a series of models to test how sensitive our response variables (namely steepness, linearity and DCI) were to those properties of the dominance matrices that are linked to sampling effort and demography – namely the matrix sparseness (proportion of pairs with no observations), the number of interactions, study duration and the number of individuals composing the matrix. In addition, we examined whether the hierarchy measures varied depending on the conditions in which the animals were observed (i.e., wild or captivity). Each predictor was evaluated in a separate model to estimate its overall association with variation in hierarchy steepness, linearity and DCI. Finally, in order to examine whether the properties of hierarchies depend on the type of behaviour used to establish dominance matrices, we tested whether hierarchies based on signals differed in steepness from those based on aggressive acts. Steepness was modelled as a function of the type of behaviour used to define dominance interactions, by including behaviour type as a categorical fixed effect. An equivalent model was fitted using hierarchy linearity as the response variable.

For the second objective, to test whether hierarchy measures are correlated across species, we fitted a model in which steepness (values bounded between 0 and 1), was modelled using a Beta distribution. We first put hierarchy linearity as a linear predictor fixed effect while controlling for matrix sparseness and the phylogenetic non-independence among species, reflected as a covariance matrix based on the phylogeny (closely related species exhibit more similar hierarchy characteristics than distantly related ones). For comparison, we also fitted a model without phylogenetic relationships, to evaluate whether the relationship between steepness and linearity would. Then, we repeated the analysis with DCI as the predictor of steepness, using the same controls. Finally, we modelled DCI as a function of linearity, again accounting for phylogeny, to test whether these two hierarchy measures were correlated across species.

For the third objective, we built multiple models to assess whether hierarchy measures differ between males and females. We fitted a model with steepness as the response variable (Beta distribution), sex as a fixed effect, and species-sex combination as a random effect to account for multiple observations per species and sex (i.e., steepness values from the same species-sex combination are expected to be more similar than across combinations). To account for phylogenetic relatedness, we fitted two separate models, one with male steepness and one with female steepness as the response variable (Beta distribution), with no fixed effects other than the intercept. Phylogenetic relatedness among species was included as a random effect, modelled as a covariance matrix derived from the phylogenetic distance matrix (37). Species identity was additionally included as a random effect to account for residual variation and multiple observations per species. Male and female hierarchies were modelled separately because the social systems of the two sexes could possibly evolve independently within a species. We subsequently contrasted the posterior distributions of the estimated means for each sex to assess whether steepness differed between males and females. We repeated the same framework for linearity and DCI. Even if male and female steepness values would not differ in these analyses across species, our sample could mask a situation where male and female steepness differ within species but in inconsistent direction across species. In other words, steepness could be higher in females of some species, and in males of other species, which in our sample could result in an overall lack of sex differences across species. In order to investigate this possibility, we examined the differences in hierarchy steepness between the sexes *within species,* selecting species for which data were available for both males and females. Specifically, we (1) calculated mean sex differences in steepness (Male_Steepness – Female_Steepness) for each qualifying species; (2) generated a null distribution of differences between female and male steepness values by randomly pairing values across species; and (3) fitted a Bayesian linear model to test whether within-species differences deviated from this cross-species null distribution. We performed this comparison by fitting a model in which the sex difference in steepness was modelled as a function of whether the sex-difference was calculated within the same species or across species (by fitting a binary fixed effect “within-species / between-species”). If the within-species indicator had a clear effect, this would suggest that sex differences within species deviate from those expected based on comparisons across species.

For the fourth objective, we tested whether potential differences in steepness and linearity between male and female hierarchies could be explained by structural properties of the matrices. We fitted models in which steepness (response variable) was modelled as a function of sex (fixed effect) and either the standardized number of individuals included in the hierarchy (fixed effect) or the log-transformed number of recorded interactions (fixed effect). We further tested whether sex differences in linearity were associated with group size by fitting a Beta regression with linearity as the response variable, sex as a fixed effect, and the number of individuals as a covariate.

For the fifth objective, to test whether sex differences in steepness values may reflect sex differences in agonistic behaviour, we ran two distinct models. To test whether hierarchical properties were linked to which sex was dominant, we used the subset of hierarchies for which information on intersexual dominance was available (average percentage of intersexual agonistic interactions won by females). We fitted a joint Bayesian model in which steepness values for males and females were modelled simultaneously as separate response variables. For each sex, steepness was modelled as a function of the proportion of intersexual fights won by females and matrix sparseness. We included an interaction between sex and the proportion of intersexual fights won by females, allowing the slope of this relationship to differ between males and females. Specifically, the slope for females was estimated directly, and the slope for males was modelled as the female slope plus an additional offset parameter. The posterior distribution of the interaction term was examined to test whether the effect of intersexual dominance on steepness differed between sexes. These analyses were restricted to the subset of 141 hierarchies (across 27 species) for which data on intersexual dominance were available, including 2 cases in which females were the dominant sex (namely *Pan pansicus* and *Miopithecus talapoin*).

For the sixth objective, we tested whether dominance hierarchies in females are more likely to be based on signals whereas those in males rely more on aggressive acts. We then fitted a binomial model in which the type of interactions determining the hierarchy (display versus act, response variable) was modelled as a function of sex (fixed effect).

## 4. Results

### 1) How robust are steepness and linearity to demographic factors and sampling design?

Among the 156 dominance matrices included in the dataset (45–119, see Supplementary Table), 99 corresponded to female matrices and 57 to male matrices, spanning 38 species (Figure 1) across three major clades: *Cercopithecoidea* (n=115 matrices on 30 species, mean steepness: 0.84, range: 0.24 - 0.99), *Hominoidea* (n=23 on 4 species, mean steepness: 0.76, range: 0.49 - 0.94), and *Platyrrhini* (n=18 on 4 species, mean steepness: 0.80, range: 0.55 - 0.96) (Figure 2). Female dominance matrices were available for 34 species, while male matrices were available for 21 species. Eighteen species provided data for both sexes, 4 species were represented exclusively by male matrices, and 17 by female matrices.

**Figure 2:**
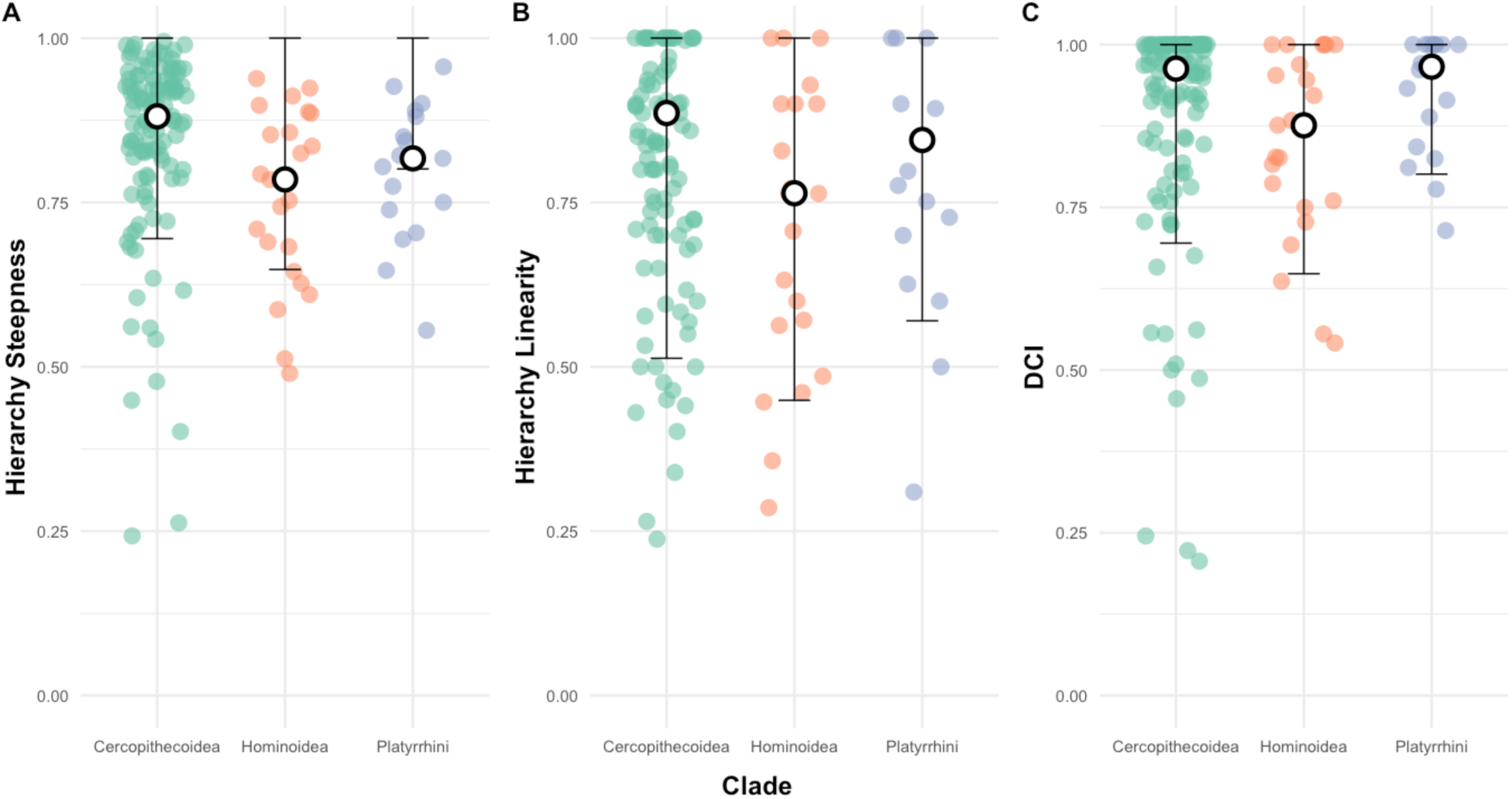
Hierarchy properties in the main primate clades. (A) Hierarchy steepness. (B) Hierarchy linearity (h’ index) and (C) Directional Consistency Index. Jittered points represent individual dominance matrices (green: Cercopithecoidea, orange: Hominoidea, blue: Platyrrhini). Central white circles indicate median values per clade; black error bars show 90% intervals.

Firstly, we assessed the robustness of our dataset by investigating whether hierarchy steepness, linearity, and directional consistency were sensitive to demographic factors and sampling design. Steepness decreased as matrix sparseness increased (posterior mean = −0.79, 89% credible interval [−1.54, −0.06]), indicating that hierarchies are less steep when fewer interactions are observed. Including study duration in the model did not alter the negative relationship between sparseness and steepness (posterior mean = −0.80, 89% credible interval [−1.54, −0.05]). Study duration itself had no effect on steepness (posterior mean = 0.00, 89% credible interval [−0.01, 0.02]). Steepness increased with the number of recorded interactions (posterior mean = 0.24, 89% credible interval [0.16, 0.33]), and the number of individuals in the group (posterior mean = 0.16, 89% credible interval [0.04, 0.28]. After accounting for the number of individuals in the group, there was no detectable difference in steepness between wild and captive populations (posterior mean difference = −0.02, 89% credible interval [−0.08, 0.04]).

Hierarchy linearity was also strongly influenced by matrix sparseness. Linearity decreased as sparseness increased (posterior mean = −0.83, 89% credible interval [−0.98, −0.69]) but was not influenced by the number of recorded interactions (posterior mean = 0.04, 89% credible interval [−0.07, 0.15]). In contrast to steepness, linearity decreased as the number of individuals increased (posterior mean = −0.48, 89% credible interval [−0.62, −0.33]). Similarly to steepness, there was no difference in linearity between wild and captive settings (posterior difference =-0.01, 89% credible interval [-0.08, 0.05]).

Directional consistency did not depend on sparseness (posterior mean = 0.11, credible interval [-0.01; 0.24]), but decreased as the number of recorded interactions increased (posterior difference =-0.14, credible interval [-0.24;-0.04]). Again, there was no difference in directional consistency between wild and captive settings (posterior difference =-0.05, credible interval [-0.11; 0.02]).

Finally, we assessed whether the steepness of the hierarchies depended on the type of behaviour used to construct the matrix, comparing matrices built from only signals (n = 16) to those built from only physical aggressive acts (n = 64). The model showed that hierarchies based on signals were slightly less steep on average than those based on physical aggression (logit difference =-0.15, 89% credible interval [-0.57, 0.27]; see Figure 3A). However, the credibility interval crossed zero, indicating no consistent effect of behaviour type on the steepness of the hierarchy. Similarly, linearity did not differ by type of behaviour (logit difference =-0.15, 89% credible interval [-0.61, 0.33], Figure 3B).

**Figure 3:**
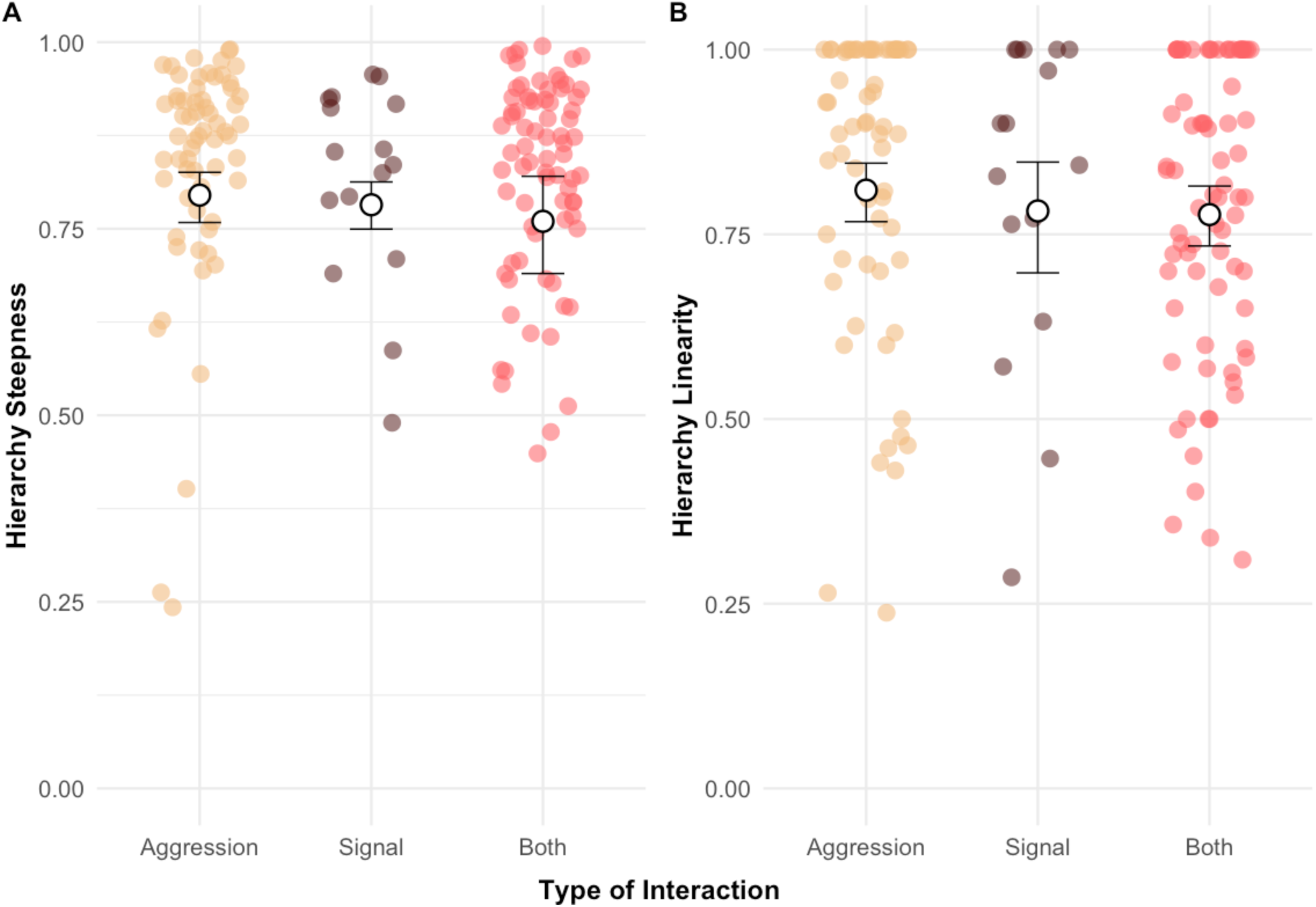
Effect of the type of behaviour used to construct the dominance matrix on steepness (A) and linearity (B). Shown are medians (black circles filled with white and its 89% CI (error bars). The sand-colored points represent matrices constructed solely from aggression, the brown points represent those constructed solely from signals, and the pink points represent those constructed from both.

### 2) Do properties of hierarchies co-vary as systems?

Across species, hierarchy steepness increased with hierarchy linearity, albeit with a lot of noise (Figure 4A). This relationship remained positive after accounting for phylogenetic relatedness and matrix sparseness (posterior mean = 1.31, 89% credible interval [0.59, 2.07]). Similarly, we found a positive association between the steepness of the hierarchy and the DCI (Figure 4B) (posterior mean = 2.85, 89% credible interval [2.18, 3.52]). However, the DCI did not increase with hierarchy linearity (posterior mean = 0.62 credible interval [0.00, 1.23]).

**Figure 4:**
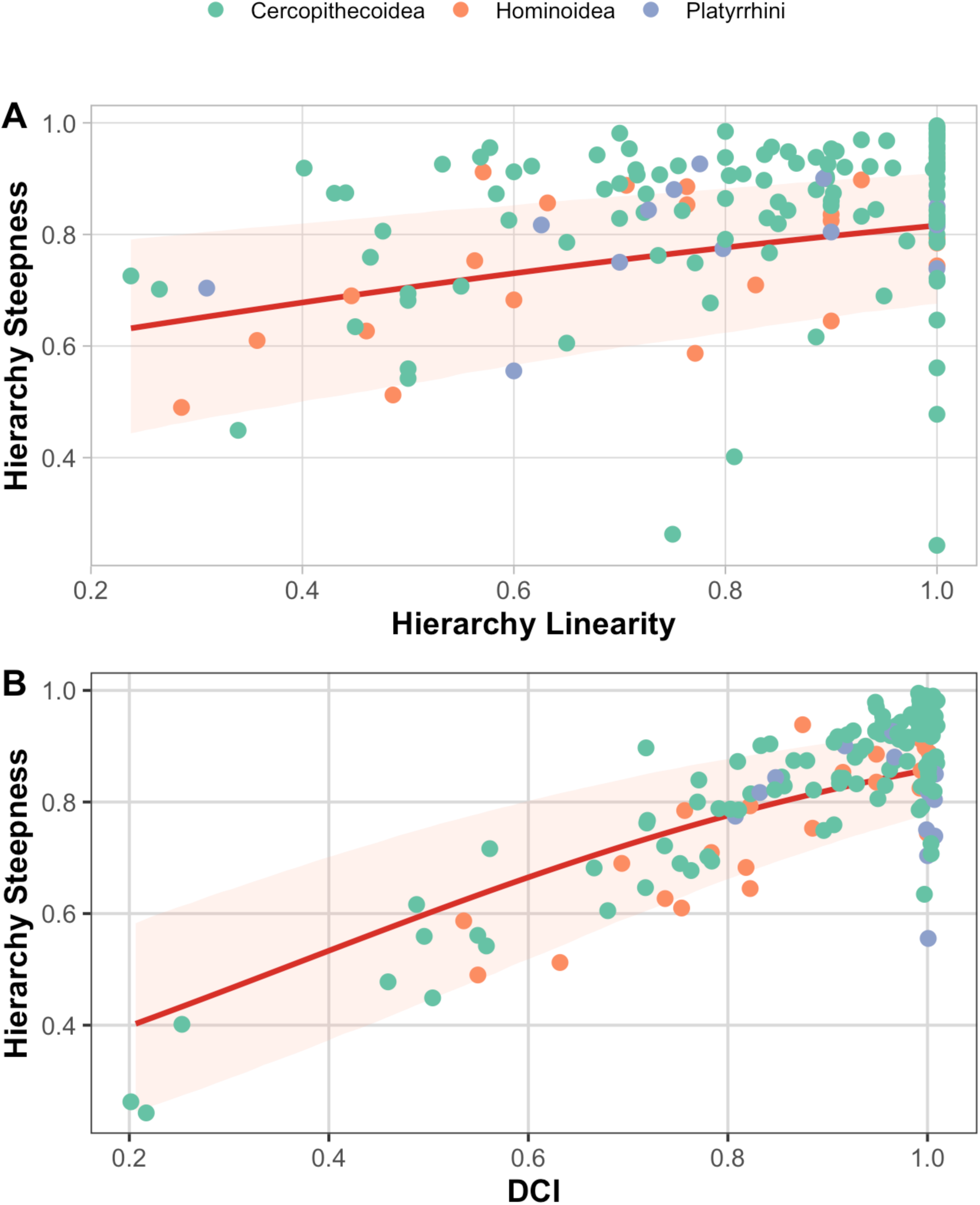
The relationship between (A) hierarchical steepness and hierarchical linearity and between (B) hierarchical steepness and the Directional Consistency Index in haplorrhine primates. The regression line is obtained from the model, and the 89% credible interval is shown. The colour of the dots indicates the clade (see legend).

### 3) Do hierarchies differ between the sexes?

Female hierarchies (n = 99) had a mean steepness of 0.84 (SD ± 0.14, range [0.24–0.99]) while male hierarchies (n = 57) had a mean steepness of 0.81 (SD ± 0.15, range [0.26–1.00]). Females thus exhibited slightly higher steepness values than males, with comparable variance. A multilevel model, accounting for repeated sampling, revealed that the estimated average steepness of female hierarchies was 0.83 (89% CI: 0.79–0.86) across species, compared to 0.82 (89% CI: 0.78–0.86) for males across species. The estimated difference (Δ males–females =-0.01, 89% CI [-0.06, 0.04]) overlapped zero, confirming the absence of a consistent difference between the sexes after controlling for repeated observations from the same species (Figure 5A).

**Figure 5:**
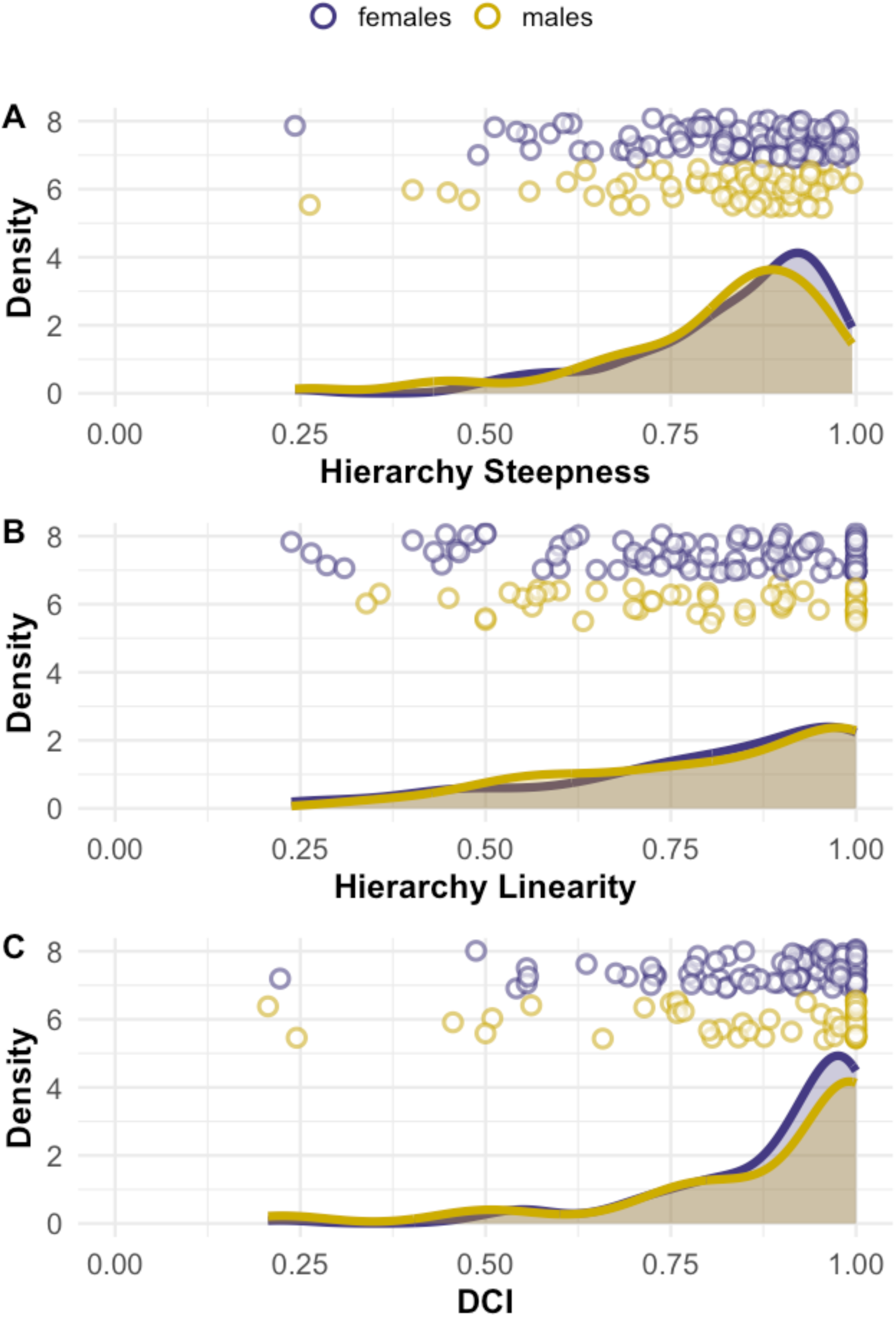
Distribution of steepness (A), linearity (B) and DCI (C) values, expressed as relative density of their occurrence for each sex. The circles represent the data points. Males are shown in gold and females in purple.

We detected a phylogenetic signal in female hierarchy steepness across species. Pagel’s λ was relatively high and significantly different from zero (λ = 0.81, p = 0.010), indicating that closely related species tended to exhibit similar steepness values. In contrast, Blomberg’s K suggested a weaker phylogenetic structure (K = 0.37) which was not statistically significant (p = 0.066). This indicates that, while closely related species tend to have similar values (i.e. species of the same Genus tend to be similar), the signal is lost when looking at more distant phylogenetic relationships (i.e. closely related Families are not more similar than more distantly related Families). In contrast to females, male hierarchy steepness did not exhibit a detectable phylogenetic signal. Pagel’s λ was effectively zero and not significantly different from zero (λ < 0.001, p = 1), indicating no covariance of steepness values among closely related species. Blomberg’s K similarly suggested a weak and non-significant phylogenetic structure (K = 0.42, p = 0.062).

A model incorporating phylogenetic covariances (normalised cophenetic distances) confirmed that there was no difference in steepness between the sexes (male–female difference = –0.08, 89% CI [–0.28, 0.13], Figure 5A). In addition, for those species for which we have observations for both sexes, sex differences in steepness within species (mean =-0.001, n = 17 species) did not differ from the differences expected by randomly pairing males and females across species (b =-0.01, 95% CI [-0.09, 0.06]).

In the raw data, male hierarchies had an average linearity of 0.81 (SD ± 0.19, range [0.31–1.00]) and female hierarchies similarly had a linearity of 0.81 (SD ± 0.20, range [0.24– 1.00]). A multilevel model estimated posterior means of 0.82 (89% CI: 0.78–0.85) for females and 0.83 (89% CI: 0.79–0.87) for males, so again the sex difference was small and uncertain (Δ males–females = 0.01, 89% CI: −0.03–0.06). Phylogenetic signal analyses confirmed minimal evolutionary structure in hierarchical linearity for both sexes. Female linearity values showed no phylogenetic signal (λ = 7.53×10⁻⁵, P = 1.0; K = 0.12, P = 0.99), whereas males exhibited a marginally stronger, albeit still non-significant, signal (λ = 7.70×10⁻⁵, P = 1.0; K = 0.35, P = 0.15). These results suggest that linearity evolves largely independently of shared ancestry in haplorhines. Phylogenetic Gaussian process models incorporating phylogenetic covariance confirmed that there is no consistent sex difference in linearity (male–female difference = – 0.08, 89% CI [–0.28, 0.13], Figure 4B).

Similarly to the other metrics, the average DCI value was high, 0.88 in males (SD ± 0.19, range [0.21-1.00]) and 0.89 in females (SD ± 0.14, range [0.22-1.00] (Figure 5C). A model based on the same design as for the other two metrics (see above) showed no sex differences (females posterior mean = 0.88, 89% CI: 0.85-0.91; male posterior mean = 0.91, 89% CI: 0.87-0.94). Among females, the value of λ was relatively high (λ = 0.779), but the result was on the margin of significance (p = 0.051). The Blomberg coefficient is low (K = 0.288) and not significant (p = 0.232). In males, λ was high (λ=0.993) but not statistically significant (p = 0.188), and K showed only a marginal trend (K = 0.474, p = 0.06). As above, phylogenetic Gaussian process models incorporating phylogenetic covariance confirmed that there is no consistent sex difference in DCI (male–female difference =-0.06, 89% CI [–0.27, 0.15], Figure 5C).

### 4) Are sex differences linked to the structural properties of their hierarchies?

Given that the number of interactions affects the steepness (see above), and primate groups usually have fewer females than males (121), we fitted an additional model that accounts for the number of interactions. Here, the difference in steepness between the sexes was, if anything, even further reduced (male-female difference = +0.01, 89% CI [-0.05, 0.07]). We then applied an analogous approach to hierarchy linearity, testing whether sex differences in linearity were associated with group size. Linearity was modelled as a function of sex and the number of individuals in the group. The estimated posterior mean was 0.81 for females and 0.79 for males, with a mean difference of −0.02 in favour of females. However, the credible interval for this difference included zero [−0.07; 0.03], indicating that there was no credible difference between the sexes in terms of linearity when the number of individuals is taken into account.

### 5) Are hierarchical properties linked to which sex is dominant?

In the subset of 141 hierarchies with matching data on intersexual dominance (here indexed by the average percentage of female wins), a model controlling for matrix sparseness showed that neither the proportion of female wins (log-transformed) nor its interaction with sex predicted steepness in either sex (female slope = 0.15 [-0.02, 0.32]; male offset =-0.16 [-0.46, 0.13]).

### 6) Do females and males use different behaviours to establish hierarchies?

Finally, regarding the behavioural foundations of sex hierarchies, 13 female hierarchies (8.3%) and 3 male hierarchies (1.9%) were based exclusively on signals, compared to 86 female hierarchies (55.1%) and 54 male hierarchies (34.6%) that involved at least one component of aggressive behaviour. A logistic model estimated the probability of a signal-based hierarchy at 15.0% [89% CI: 10.0–21.0%] among females, compared with 9.0% [5.0–15.0%] among males. The difference (males – females = −6.0% [−13.0, +2.0%]) overlapped with zero, indicating that even though there is a difference, there is no consistent sex difference in how hierarchies are established.

## 5. Discussion

Steepness, linearity, and directional consistency covary across species (species with steeper hierarchies also tend to have more linear, ordered ones), suggesting that these measures capture coherent aspects of hierarchy structure rather than independent properties. However, these measures of the structures of dominance hierarchies did not show the expected sex differences. Our finding likely results both from methodological and biological limitations in what steepness and ordering can reflect about social structure. All three measures were strongly affected by sampling design. Linearity and steepness decreased as matrices became sparser. Steepness increased with the number of recorded interactions and the number of individuals in the group, but the number of interactions had no detectable effect on linearity. Directional consistency declined as the number of recorded interactions increases. For both sexes, the structure of their hierarchies did not appear to depend on inter-sexual dominance, nor on whether dominance was established through signals or aggressive acts.

### 1) Covariation among steepness, linearity and DCI, and methodological constraints

Our results suggest that steepness, linearity, and directional consistency are not independent properties, but rather interrelated measurements reflecting how agonistic interactions occur among individuals. Across primate species, when dominance relations are more asymmetric, they are also more ordered. This concept is found in the pioneering studies on primate socioecology, which describe egalitarian societies as having a weakly linear and shallow hierarchy, and despotic societies as having a steep and ordered hierarchy (13,15). For example, rhesus macaques (*Macaca mulatta*) exhibit steep and highly linear hierarchies, while the more tolerant Tonkean macaques (*Macaca tonkeana*) are often described as having relaxed dominance relationships and lower hierarchical steepness. While the literature generally focuses on examples at these extremes (16,18,122,123), our findings suggest that these measures co-vary across species along a gradient of order. Steepness and directional consistency are particularly closely linked because they are both based on relative measures of the wins and losses among dyads. The covariation between steepness and linearity is structural i.e., imposed by their definition, because a supposedly lower-ranking individuals winning an encounter against a supposedly higher-ranking individual (i.e. A winning against B, B against C, but C against A) leads to a non-transitive hierarchy, which alters the social rank ordering and reduces steepness - the slope of the relationship between dominance and rank. Hence, hierarchies with low linearity are expected to show low steepness. However, this covariation by construction is not total, because a hierarchy can be linear without necessarily being very steep or directionally consistent. Individuals can be ordered in rank, with varying degrees of the steepness of the slope between dominance and rank. Indeed, de Vries et al. referred to this pattern to support their argument “that linearity and steepness measure two different characteristics of a dominance hierarchy” (17). However, they did not present hierarchies displaying the opposite pattern (i.e. high steepness with low linearity), and we do not observe these in our data (Figure 4) because of the limitations in non-transitive hierarchies. This suggests that steepness and directional consistency may capture aspects of hierarchy structure that are less directly constrained by the transitivity of dyadic outcomes than linearity, which appears more tightly linked to the underlying data structure (see also Koenig et al. 2013 (25)).

More broadly, very few hierarchies in our dataset had steepness, directional consistency or linearity values below 0.5, indicating that when dominance hierarchies can be identified, they typically exhibit at least a moderate degree of order (see also (124)). A straightforward explanation is that individuals within groups differ in competitive ability (e.g. in age, size, condition or experience), such that some are consistently more likely to win than others. These phenotypic asymmetries can generate and maintain ordered dominance relationships. In addition, processes such as winner–loser effects may further reinforce existing asymmetries so that individuals that win contests become more likely to win again, and those that lose become more likely to lose again (124,125). Such experiential effects can sharpen or stabilize rank differences over time, increasing linearity, directional consistency and steepness (126,127). In support of these effects, we found that both steepness and directional consistency were primarily affected by the number of interactions that occurred, independent of sparseness, study duration and the number of individuals in the group. This indicates that hierarchies are more ordered when individuals interact more frequently. This argument is further supported by our observation that the hierarchy structure remains the same regardless of how hierarchical dominance relationships are expressed (through acts or signals, aggressiveness or submission), the hierarchy, the structure appears the same.

Our results indicate that Bayesian estimates of steepness, the directional consistency index and the linearity of the hierarchy are affected by demographic and sampling properties of the data, such as group size, the number of recorded interactions and matrix sparseness, to a greater extent than is generally acknowledged (23). In other words, how many individuals are included and how intensively they are observed both tend to generate more recorded dominance interactions in addition to individuals actually showing more frequent aggression, and the three metrics vary substantially as a function of this interaction. In practice, this means that variation in steepness, linearity and DCI is affected by group size and sampling effort, rather than solely capturing variation in hierarchical behaviour. In addition to this sensitivity, steepness and linearity cannot differentiate a genuine absence of aggression among pairs of individuals - which indicate low linearity and low steepness – with missing observations (25). While the Directional Consistency Index does not solve the problem of ties and missing interactions, it is affected by them in a different way than steepness or linearity. Indeed, dyads with no recorded interactions—whether because individuals never interact aggressively or because interactions were not observed—do not enter the DCI calculation and therefore do not change its value. By contrast, dyads with observed interactions in both directions contribute to lower DCI values, reflecting genuinely bidirectional relationships. Unlike steepness, DCI decreases with an increasing number of observations. This effect likely indicates a higher chance of observing an aggressive act that occurs against the primary direction, as well as the fact that the DCI is a ratio, with the number of interactions as the denominator (24). Overall, it currently remains difficult to determine when a group has been sampled sufficiently to accurately reconstruct the relationships among individuals, making it difficult to evaluate the extent to which our set of metrics reflect variation in hierarchical behaviour above and beyond variation in sampling effort or demographic properties such as group size.

Steepness and directional consistency nevertheless appear to capture some biological variation in the structure of primate groups, if only because both measures show a phylogenetic signal for females, with closely related species showing similar values (e.g. females in baboons tend to live in matrilineal societies). However, this phylogenetic signal may partly reflect similarities in socio-demographic properties among related species, such as group size and group composition, rather than exclusively differences in hierarchical behaviour (see also (128)). Previous comparisons of steepness were mostly restricted to few, closely related species and might therefore have been less affected by these biases. We did not find a phylogenetic signal for the steepness of male hierarchies, potentially reflecting that their social structure is less conserved among closely related species or alternatively that the female analysis was more powerful because the dataset included more female than male hierarchies. Indeed, primate groups typically contain more adult females than adult males, largely as a consequence of sex-specific dispersal patterns and female philopatry (4,13). In addition, our male sample may be biased against species with the most intense within-group competition. In such species, intense male–male aggression is likely to lead to the exclusion of subordinate males from the group and the formation of one-male units or groups, rather than to the formation of steep, stable hierarchies. These groups are not represented in our dataset because hierarchies only exist in multi-male groups.

### 3) There is no sex-specific bias in the structural properties of the dominance hierarchies

Our results show that the measures of hierarchical properties we used here cannot reveal the potential sex differences in competition. Two interpretations can account for the absence of sex differences in steepness, linearity and directional consistency. First, these metrics may be too sensitive to methodological factors such as group size, sampling effort, and heterogeneity in study designs to reliably capture biologically meaningful variation in dominance structure in a comparative context (see the section above). On the other hand, these metrics might reflect biological meaningful variation in behaviour, however only in behaviour that does not reflect the level of social competition. With our broad-scale comparative approach, we cannot tease the methodological and the biological limitations apart. Both probably interact to explain the variation in the measured structures of the hierarchies because we observed that accounting for the methodological factors further reduced any sex differences in steepness, linearity, and directional consistency. We also found that steepness, linearity, and directional consistency did not differ depending on the behaviour that was used to build the hierarchies. These measures therefore might simply reflect how frequently individuals interact (127) and how established their relationships are during the study period, but not the degree of competition and potential skew in power among individuals. Our study highlights the need for alternative or complementary metrics to capture variation in dominance structures, ideally those that are robust to variation in sampling and socio-demographic factors (see. e.g (129,130)).

There is evidence to suggest that, in mammals where females are dominant over males, their hierarchies rely on fewer aggressive acts but more submission signals than in the male-dominant societies. This difference in behaviour indicates that conflicts are resolved differently in groups (28). A recent comparative study classified species as ranging from strictly male-dominant to strictly female-dominant, based on the percentage of contests won by females (27). Applying this same average percentage to our species, we however found no variation in the steepness of the hierarchies of either sex in relation to the variation in the degree of female dominance. This reinforces the idea that the measures of hierarchical structures we used here do not appear to capture variation in the sources of competition or which sex dominates.

## 6. Conclusion

### Conclusion and outlook

Across the primate species and groups included in our analyses, male and female dominance hierarchies were remarkably similar in their steepness, linearity and directional consistency. This absence of sex differences suggests that, at least as captured by these metrics, the structural properties of dominance hierarchies do not differ systematically between the sexes, despite well-documented differences in the sources, mechanisms and forms of social competition. However, given that these three metrics mainly reflect sampling effort and shared patterns of behaviour, our study remains inconclusive regarding the biological meaning of the absence of sex differences in hierarchical structure. Our findings instead suggest that the problems that troubled researchers of primate dominance decades ago (e.g. (24,131) persist today. While we can rank individuals within groups, we still lack good measures that would capture meaningful variation in dominance structures across groups and species. To understand how the strength and resolution of competition and conflict differ across societies, we require different ways to capture the dimensions of power, including alternative metrics of asymmetry and inequality that are less sensitive to sampling intensity and data sparseness.

## Acknowledgments

We thank Eve Davidian, Marta Mosna, Oliver Höner, Tal Kleinhause, and Christof Neumann for helpful feedback during the completion of the study.

## Ethical Statement

Our study relied on previously published data and did not involve working directly with animals.

## Data Accessibility

All data and code required to repeat all analyses in this manuscript are currently accessible at github: https://github.com/dieterlukas/DominanceHierarchyDifferences and upon acceptance of the manuscript will be deposited at Zenodo.

## Authors’ Contributions

L.S.: data curation, formal analysis, investigation, methodology, validation, visualisation, writing – original draft, writing – review & editing, final approval

E.H.: conceptualization, formal analysis, funding acquisition, investigation, methodology, validation, writing – review & editing, final approval

D.L.: conceptualization, data curation, formal analysis, funding acquisition, investigation, methodology, validation, writing – review & editing, final approval

---

**Supplementary Table S1:**
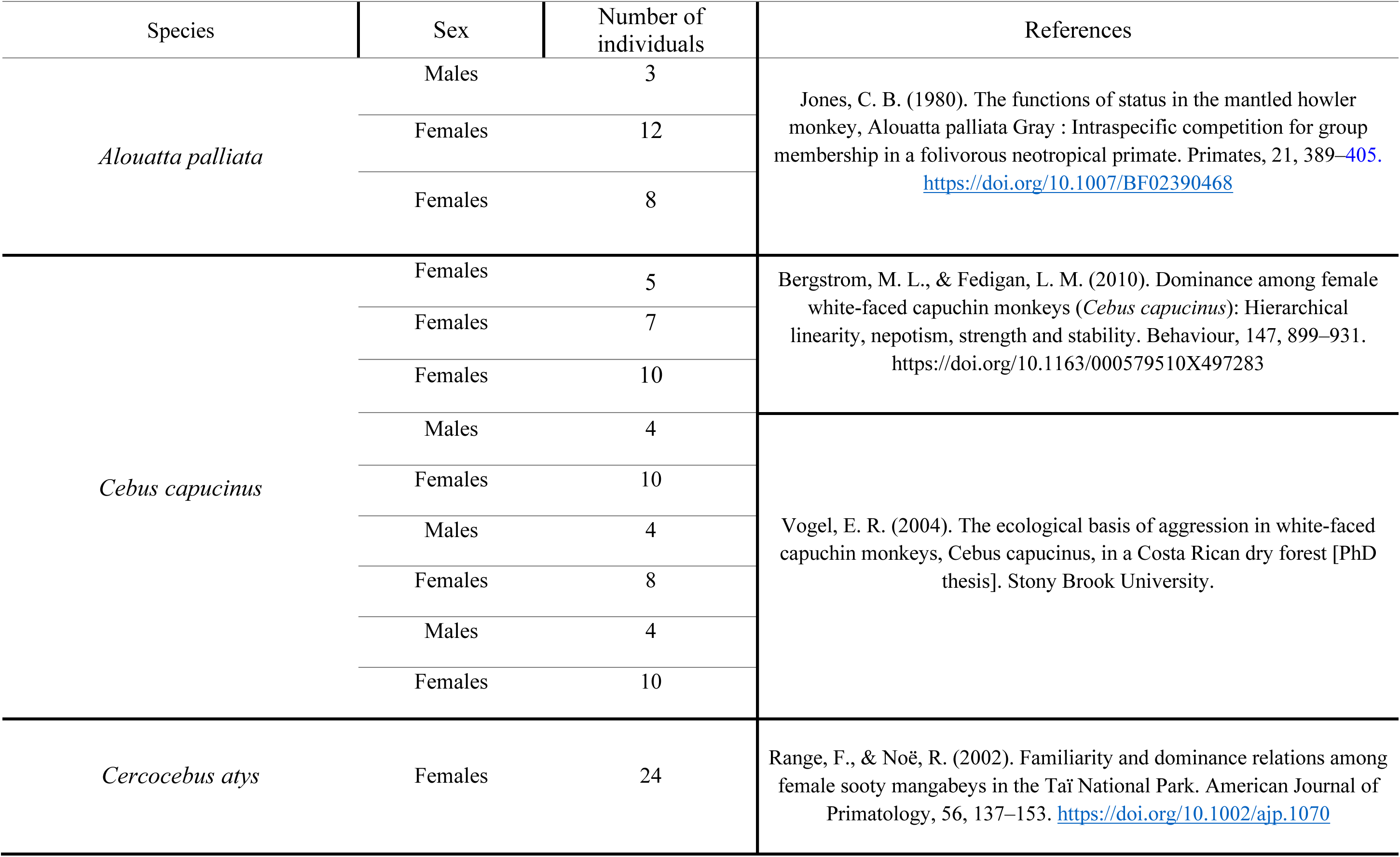
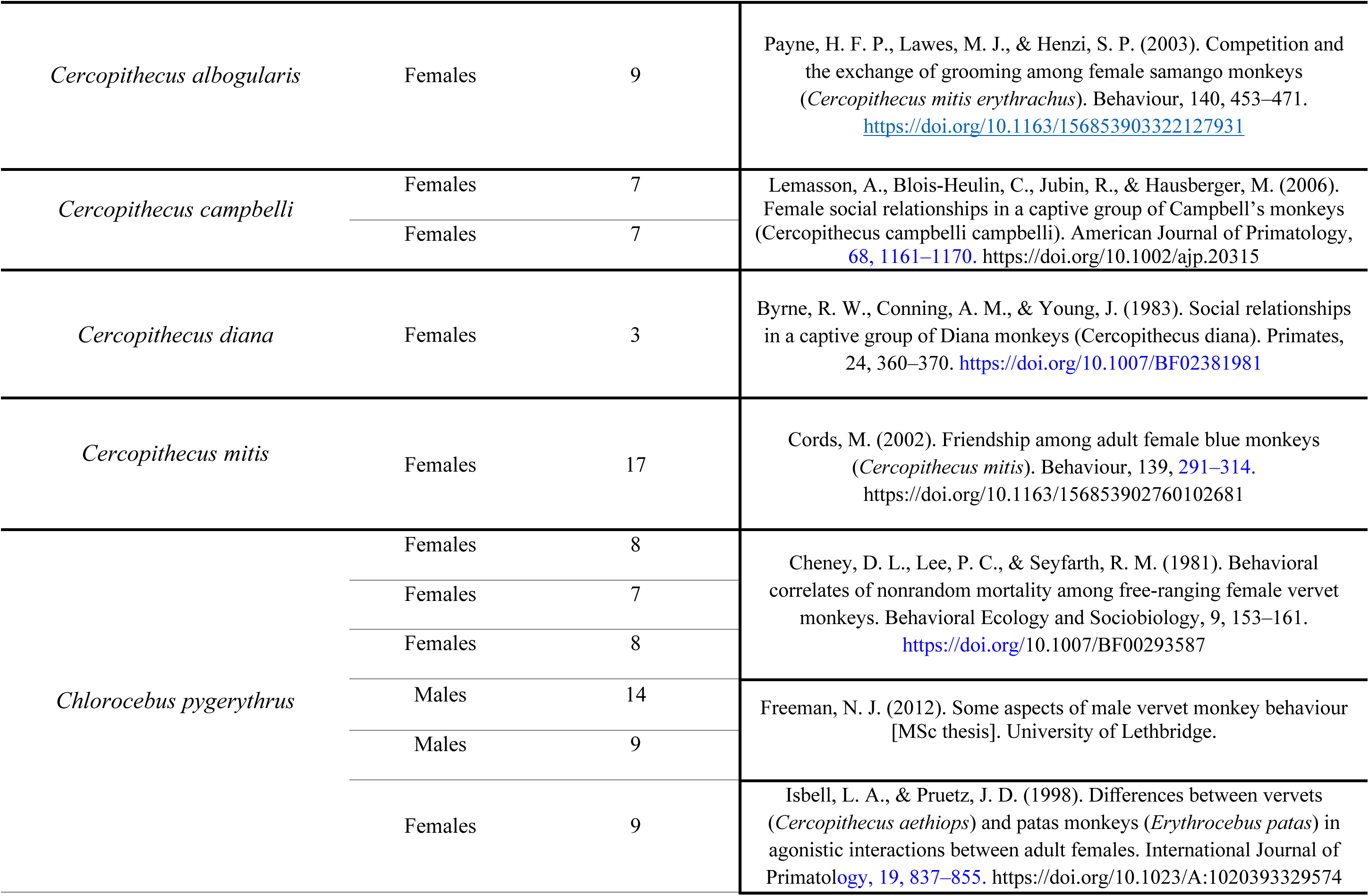
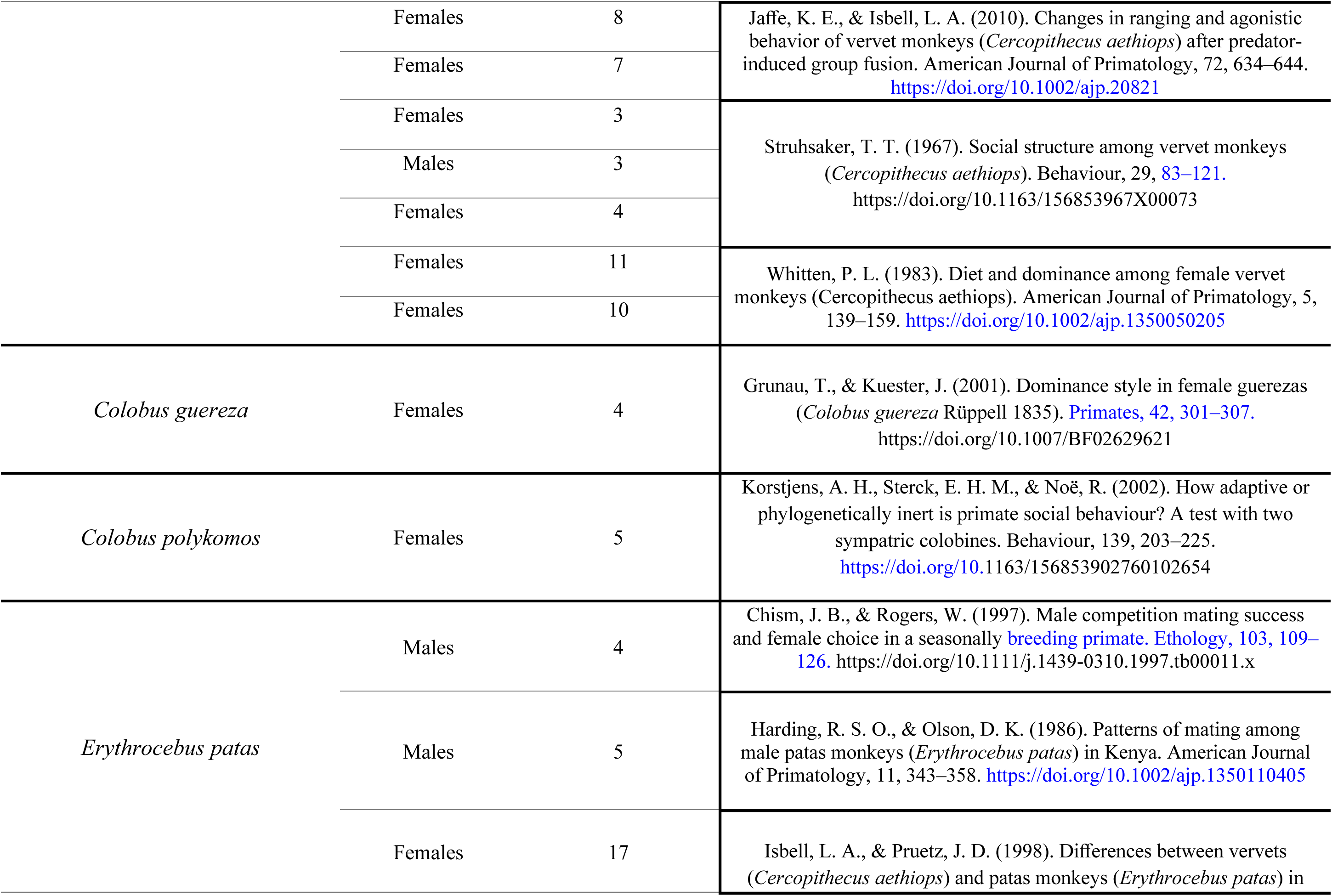
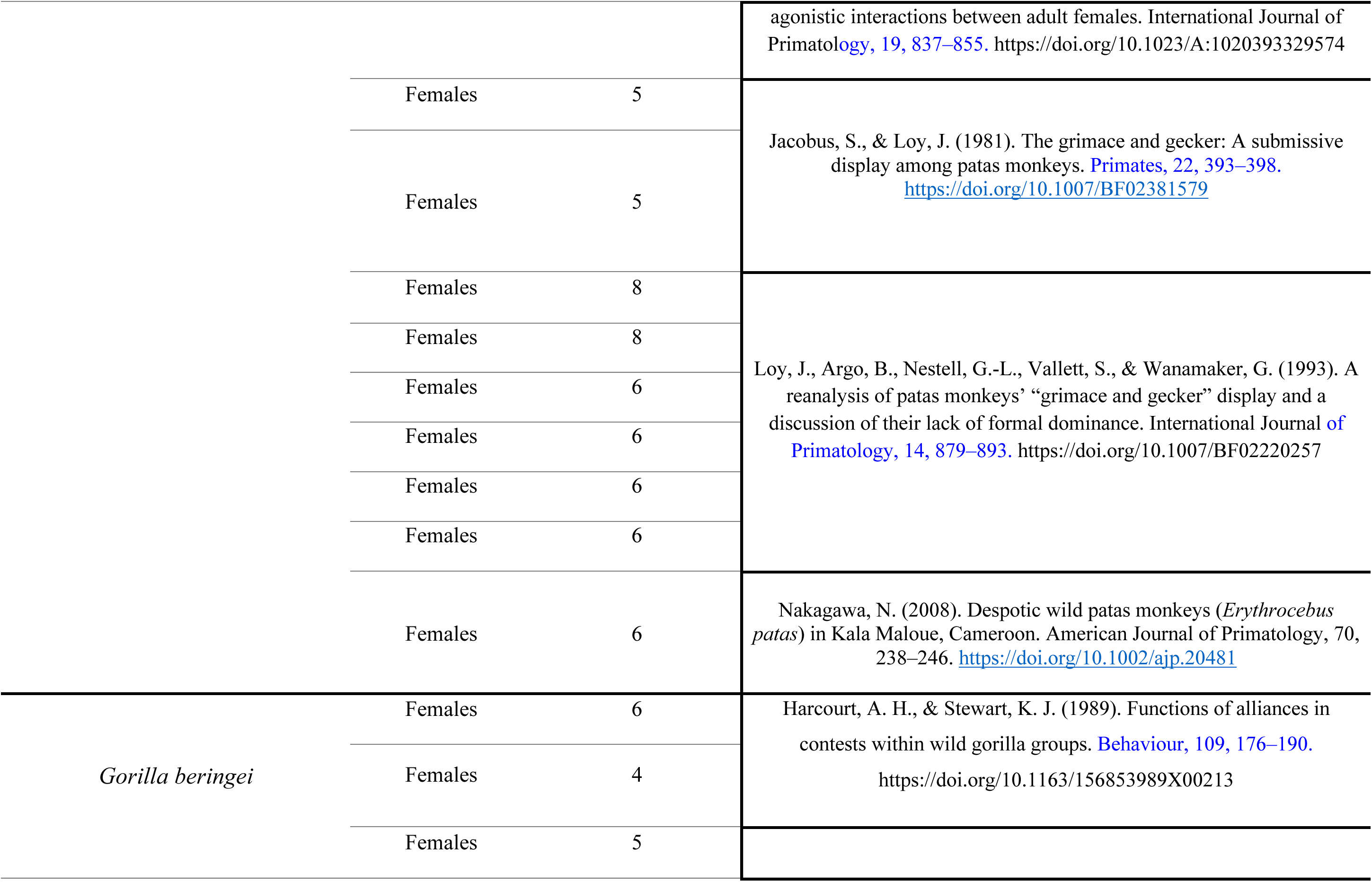
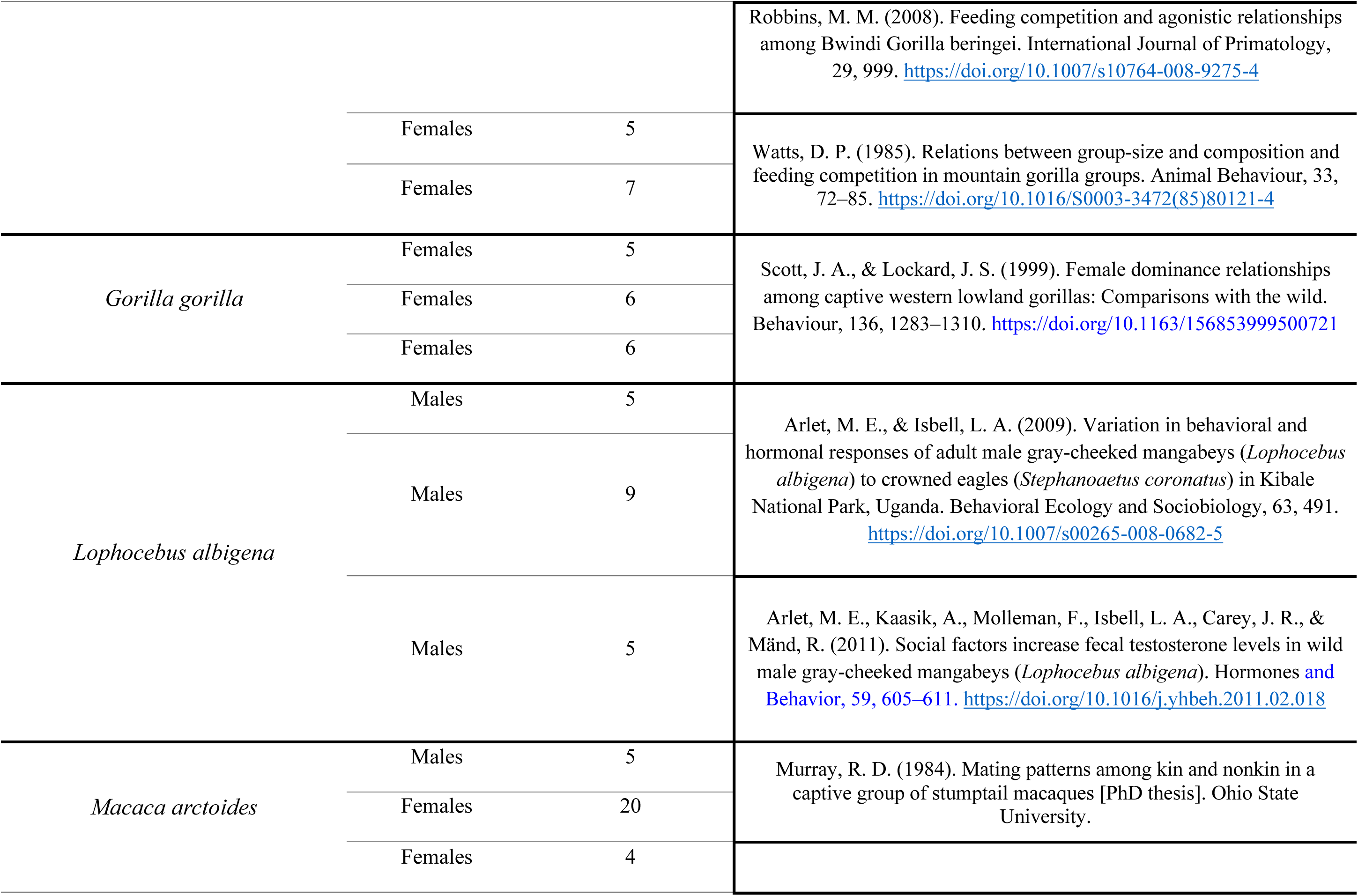
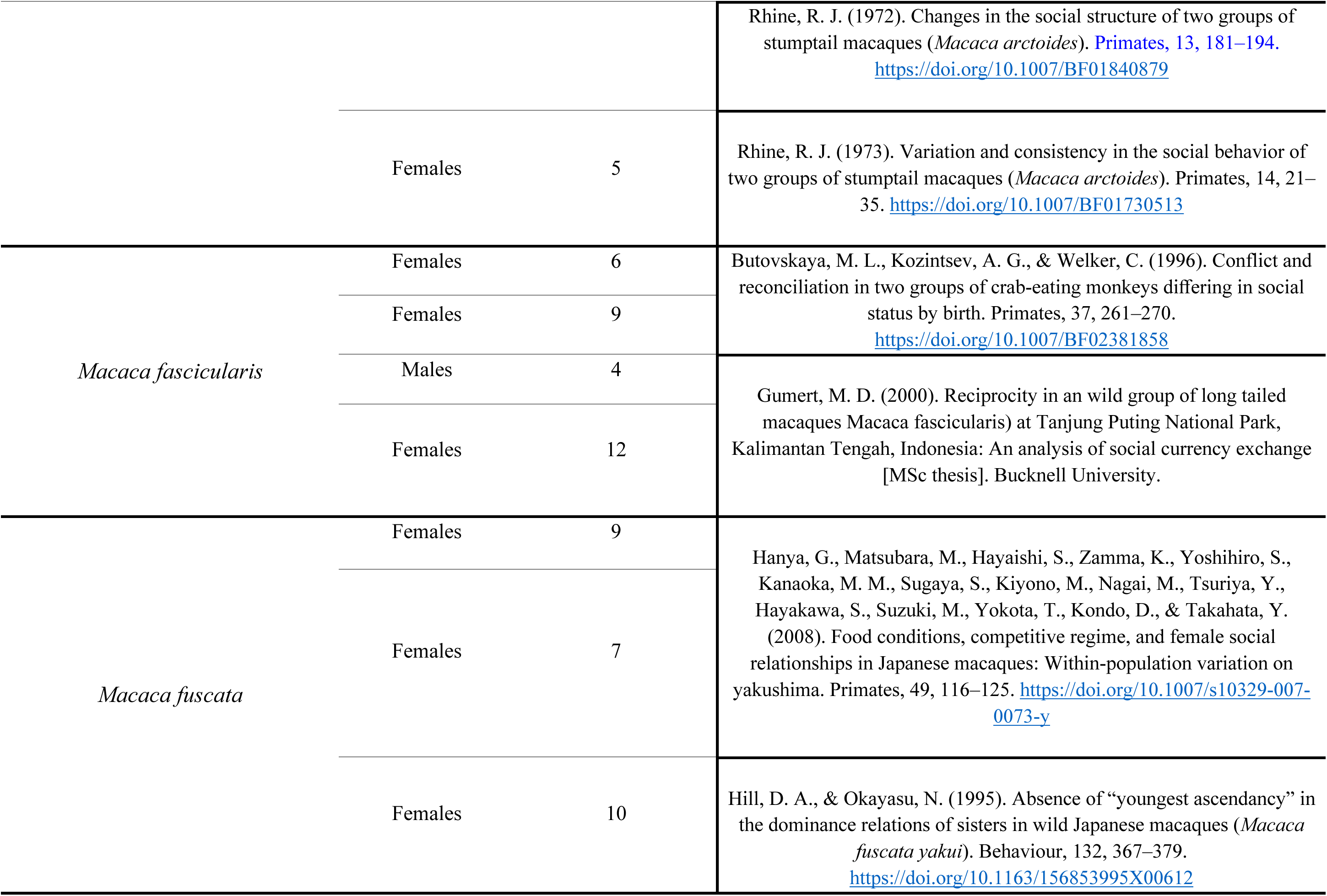
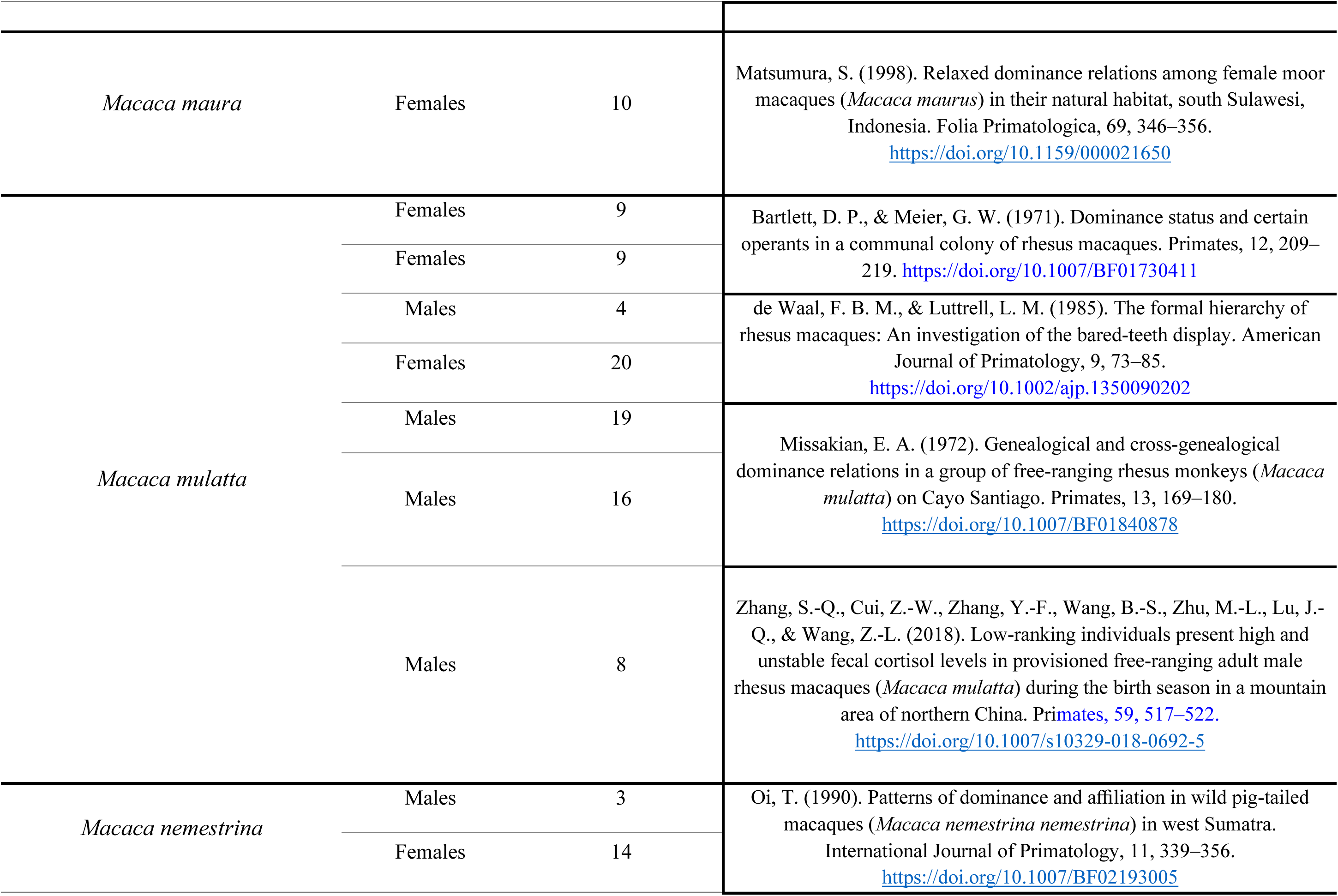
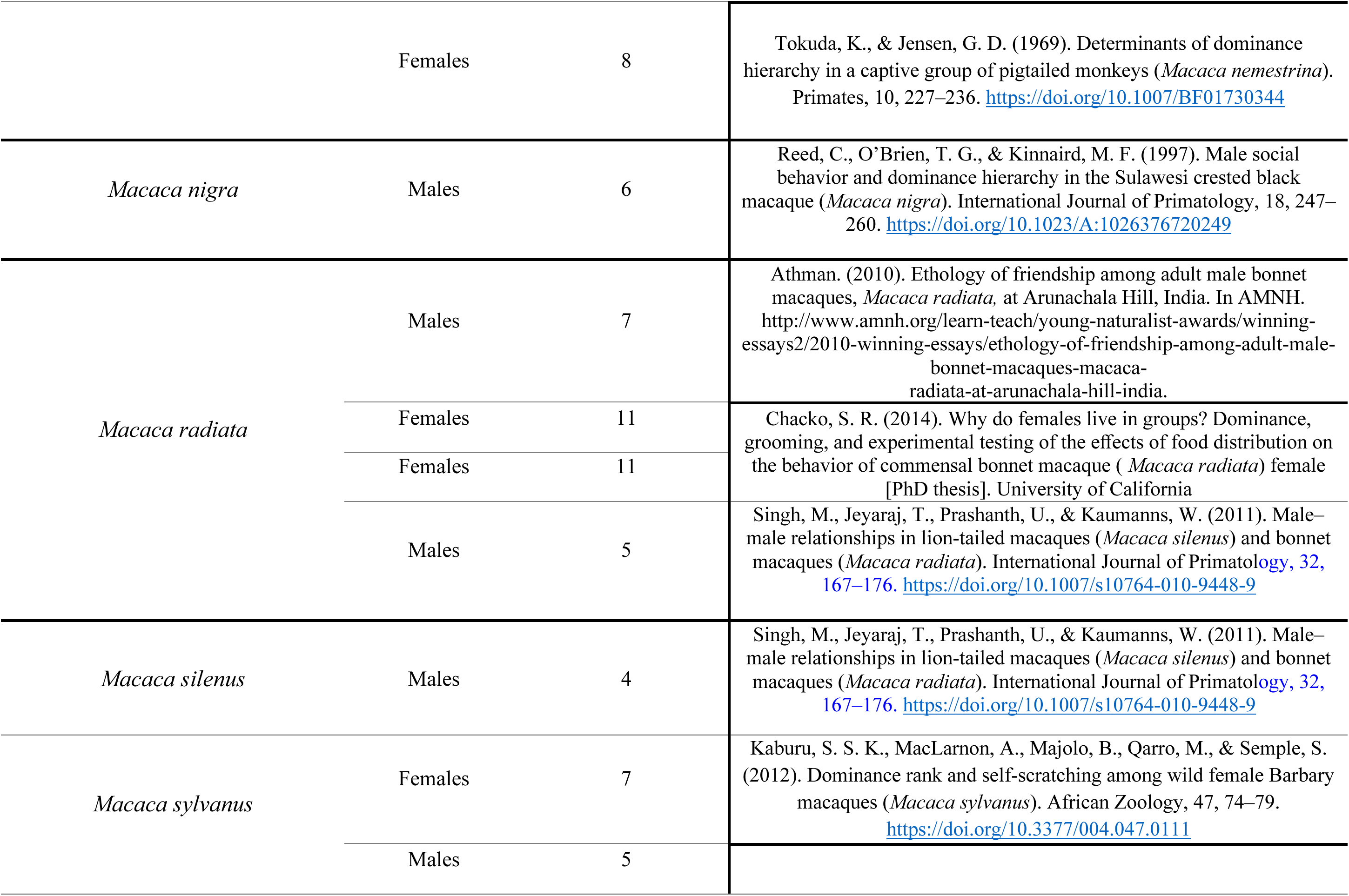
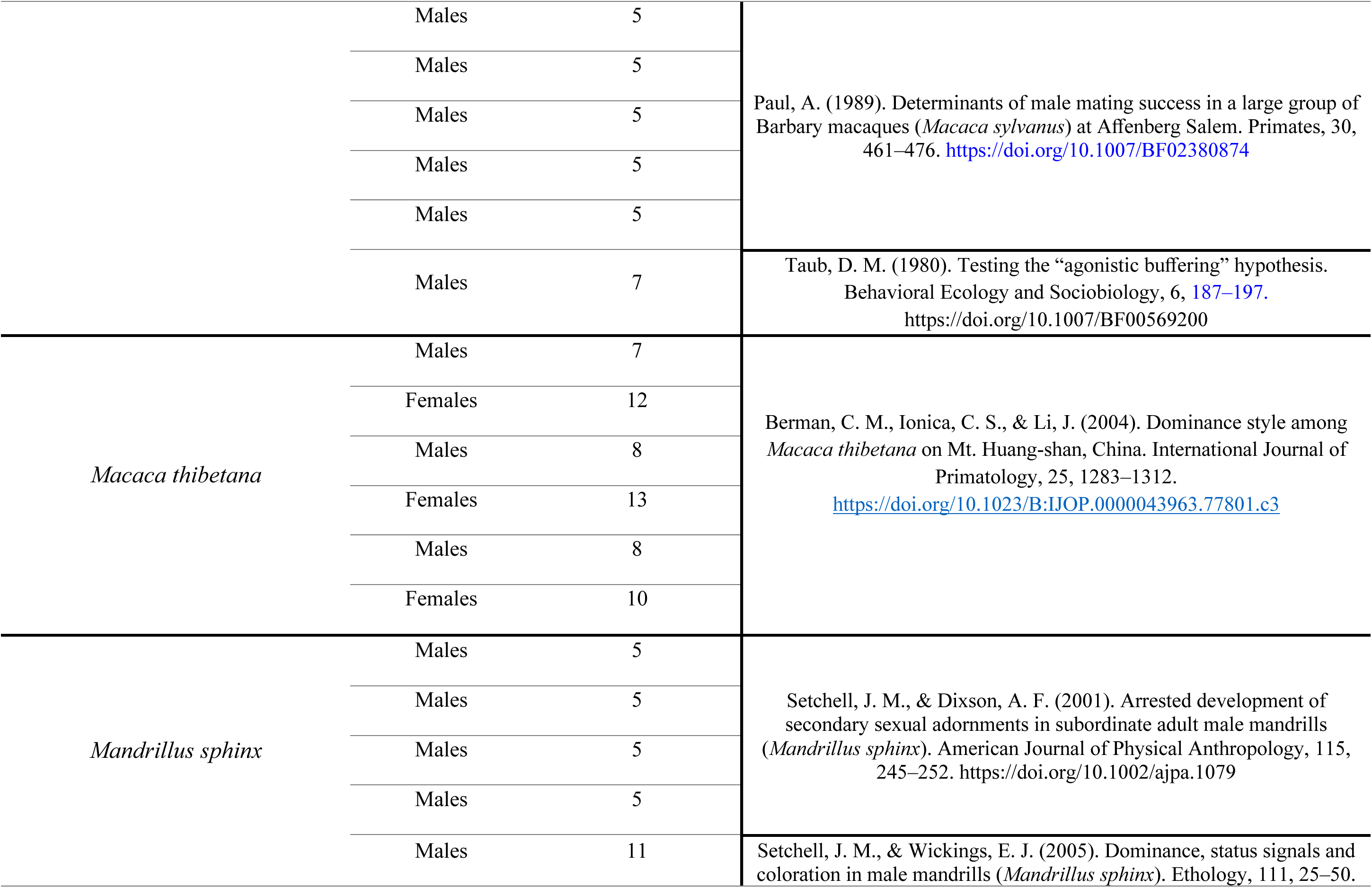
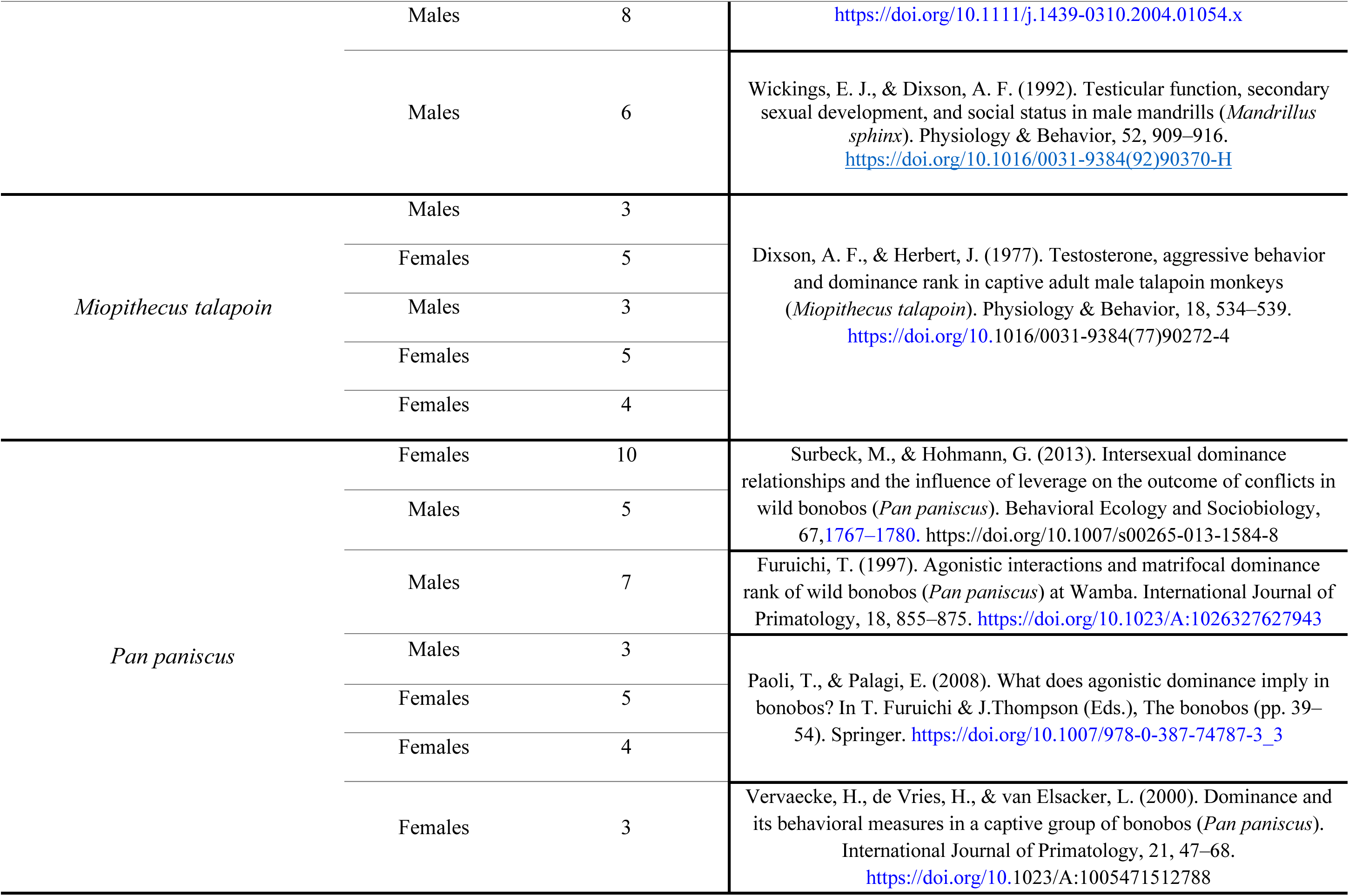
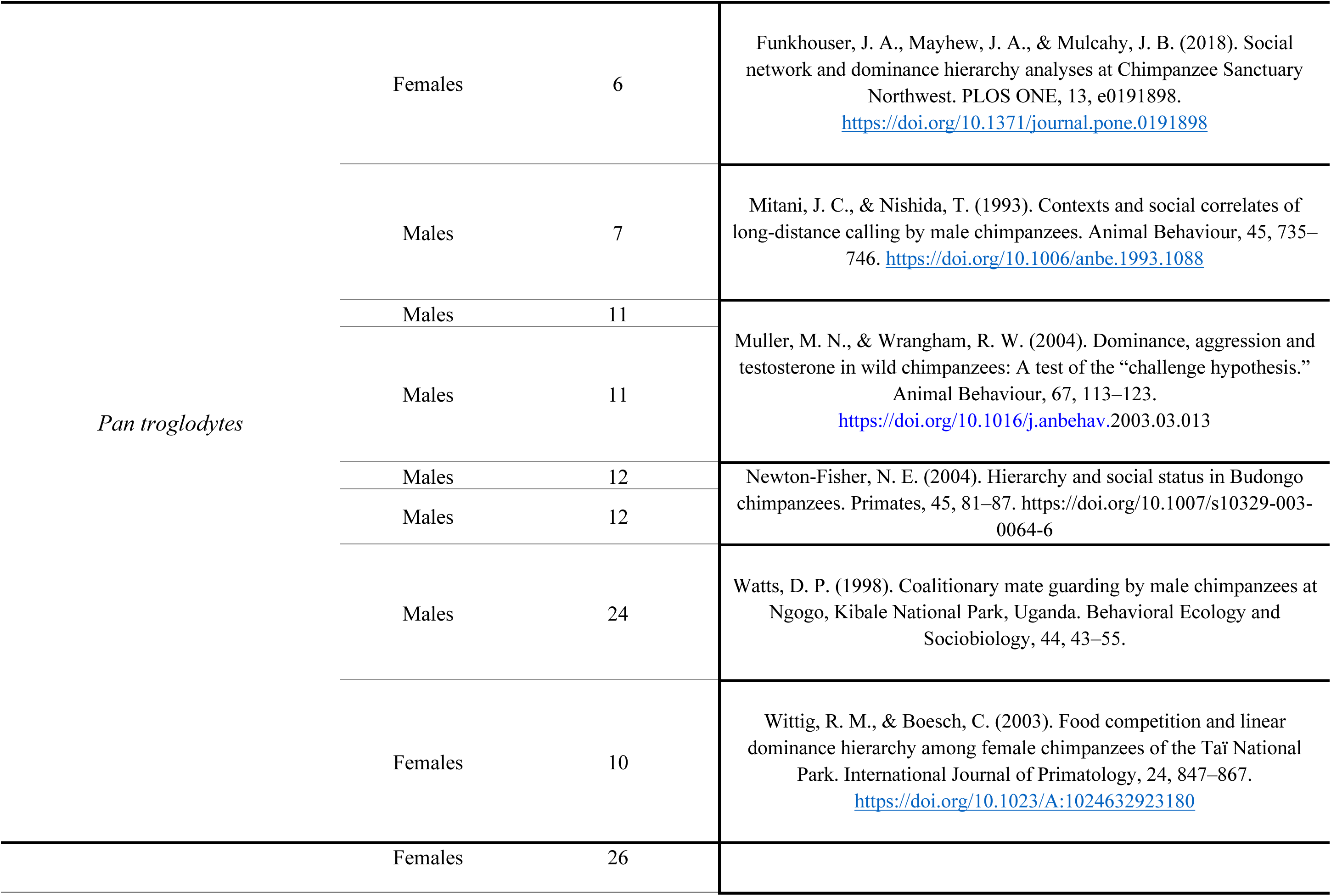
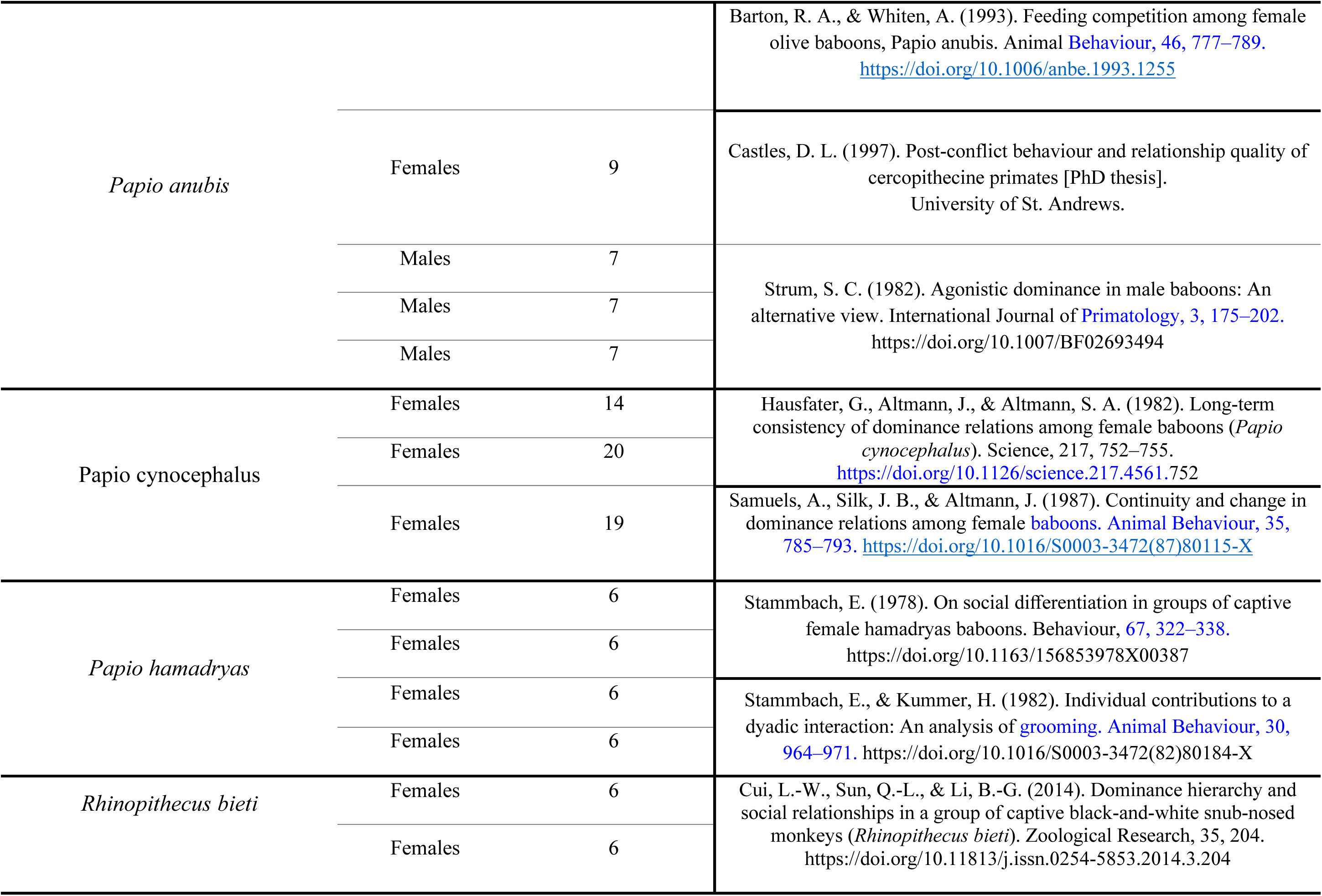
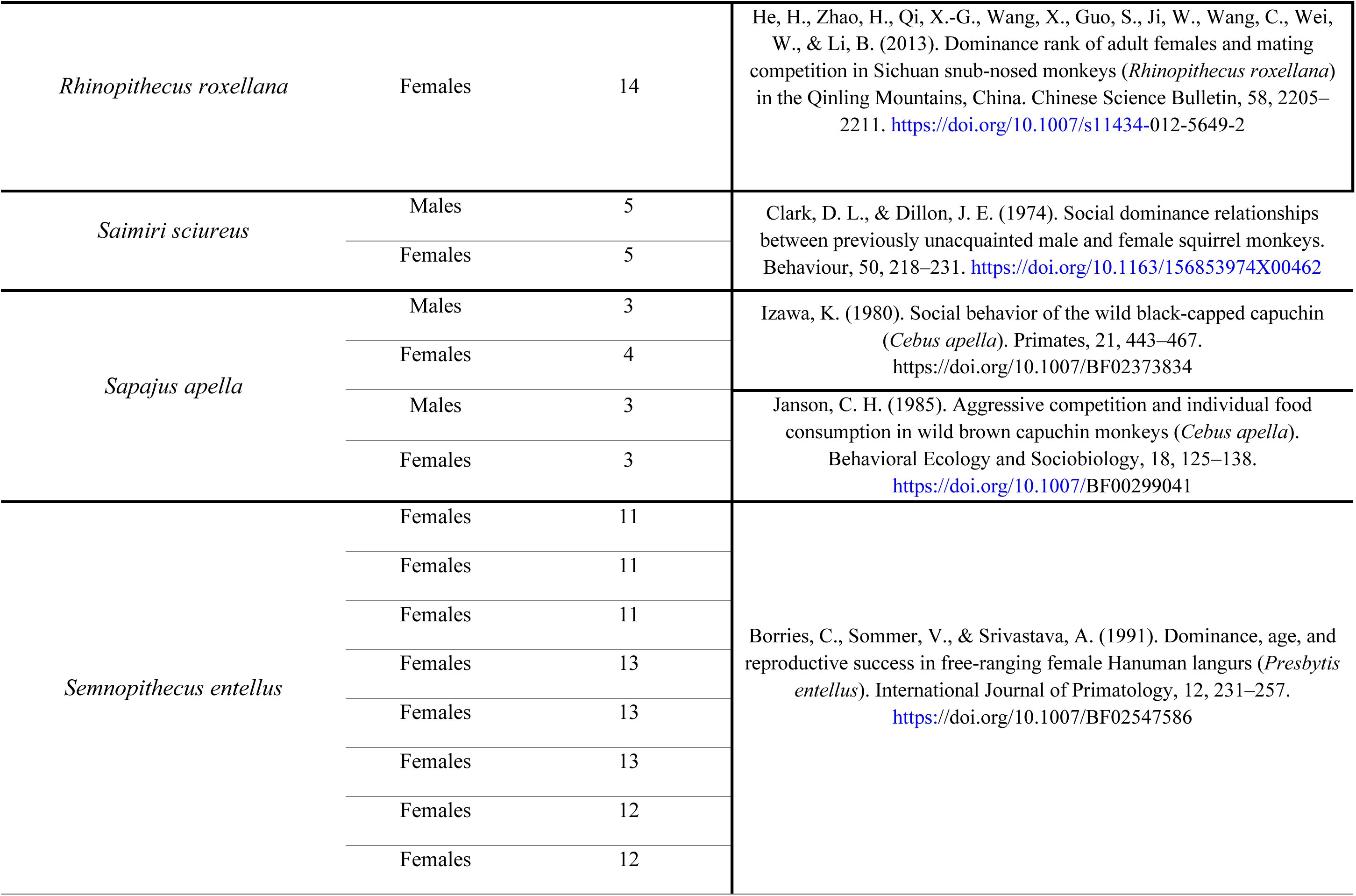
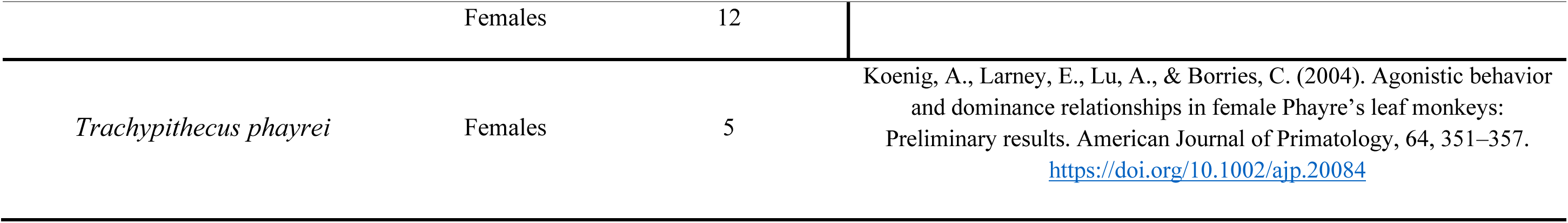
Original data with references from the EloSteepness.data database (Neumann, 2022)

